# Genome-wide Chromosome-specific Aneuploidy Engineering and Phenotypic Characterization with CRISPR-Taiji

**DOI:** 10.1101/2025.04.25.650684

**Authors:** Hugang Feng, Daqi Deng, Rashmi Dahiya, Libin Wang, Jingkun Zeng, Benjy Jek Yang Tan, Fiona Byrne, Scott T. C. Shepherd, Jiahao Wang, Sarah C. Johnson, Alison Harrod, Karen A. Lane, Annika Fendler, Anne-Laure Cattin, Zayd Tippu, Meilun Nie, Yiming Zhao, Ruijia Wang, Wei Ai, Omar Bouricha, Taja Barber, Yuliia Dovga, Yihan Xu, Weiming Shen, Zhen Sun, Hongye Wang, Jiufei Zhu, Yimeng Xu, Liani G. Devito, Lyn Healy, Eugénie S. Lim, Samuel M. O’Toole, Scott Akker, William M. Drake, Haojie Jin, Jessica A. Downs, Sarah E. McClelland, John F. X. Diffley, Peter Ly, Samra Turajlic

## Abstract

Aneuploidy, the gain or loss of chromosomes, is prevalent in both normal and disease conditions, however, experimental approaches to engineer and study aneuploidy remain limited, leaving its functional significance under-characterized. Here, we present CRISPR-Taiji (CRISPRt), an efficient method for inducing chromosome-specific mis-segregation and aneuploidy generation across all 24 human chromosomes via dead Cas9 (dCas9)-induced centromeric chromatin relaxation. Using CRISPRt with scRNA-seq, we generated the first comprehensive transcriptomic alteration landscape of nearly all autosomal aneuploidies at chromosome-arm resolution. This genotype-phenotype map provides causal evidence linking recurrent aneuploidies in clear cell renal cell carcinoma (ccRCC) to molecular and clinical phenotypes observed in patient tumors. Notably, chromosome 3(p) loss, the ccRCC initiating event, specifically drives strong interferon signaling activation, offering novel insights into ccRCC tumorigenesis and immune modulation. Overall, we establish CRISPRt as a simple, efficient and scalable approach for chromosome-specific aneuploidy engineering and characterization in preclinical models to advance aneuploidy research across diverse biological contexts.

## INTRODUCTION

Aneuploidy, the presence of whole-chromosome (whole-chr) or chromosome-arm (chr-arm) copy number gains or losses, can arise from different mechanisms, including mitotic chromosome mis-segregation. Aneuploidy in gametes or embryonic stem cells can lead to chromosomal disorders such as Down syndrome^1^. In addition, somatic aneuploidy has long been recognized as a near-ubiquitous feature in most cancer types^2^, while recent studies revealed the widespread presence of somatic aneuploidy across morphologically normal tissues^3–6^.

Like genetic mutations, distinct recurrent patterns of aneuploidies are observed across different cancer types and normal tissues, highlighting context-dependent functional significance and selection^3–7^. However, in contrast to our ability to model gene-level alterations, it has been challenging to efficiently engineer targeted aneuploidy events in relevant preclinical models for functional interrogation^8^. Consequently, despite its prevalence, the functional significance of individual aneuploidies across various biological contexts remains poorly understood, which restricts the rational development of therapeutics targeting the recurrent aneuploidies in human diseases.

To date, several approaches for generating aneuploidy have been developed. Mitotic checkpoint inhibitors, such as reversine, can efficiently generate near-random chromosome mis-segregation and aneuploidy, but lack chromosome specificity^9^. More recently, chromosome-specific aneuploidy engineering methods were developed by inducing chromosome-specific mis-segregation, including ectopic kinetochore establishment^10^, kinesin motor tethering^11^, and manipulation of the histone H3 variant centromere protein A (CENPA)^12^; however, the application of these methods is restricted to a subset of chromosomes. In addition, approaches such as microcell-mediated chromosome transfer (MMCT)^13^ and sequence-specific DNA breakage induction^14,15^ can specifically target all chromosomes, but generate only gains or losses, respectively, with limited efficiency.

Recently, the complete human centromeric DNA sequence was resolved by the telomere-to-telomere (T2T) consortium, sparking increased interest in human centromere research^16–18^. The human centromere is mainly composed of α-satellite (αSat) DNA tandem repeats, organized into higher-order repeat (HOR) structures spanning millions of bases^19^. In each human chromosome, a portion of the αSat HOR region is epigenetically defined by the incorporation of CENPA, the key nucleating component of the kinetochore, which is a protein network essential to attach chromosomes to mitotic spindle microtubules for chromosome segregation^19^. Therefore, with the newly resolved centromeric sequence, CRISPR-based tools can now directly target these centromeric regions responsible for chromosome segregation. A recently described approach, KaryoCreate, achieved chromosome-specific mis-segregation and aneuploidy generation for 10 human chromosomes by targeting mutant kinetochore component Kinetochore Scaffold 1 (KNL1^mut^) to centromeric αSat HOR regions using catalytically dead Cas9 (dCas9) for microtubule attachment disruption ^20^.

Despite remarkable advances, a simple, efficient, specific, and scalable method to engineer aneuploidy across all human chromosomes remains lacking. Here, we present such an approach using dCas9 alone, which induces chromosome-specific mis-segregation and aneuploidy generation via centromeric chromatin relaxation. Using this approach, we systematically engineered and characterized all autosomal aneuploidies in isogenic backgrounds to delineate the first genome-wide transcriptomic alteration landscape of aneuploidy. This landscape faithfully recapitulates the well-characterized interferon signaling dysregulation driven by Chr21 trisomy in Down syndrome patients. We further demonstrate its utility in establishing new causal relationships between specific recurrent aneuploidies and disease-relevant transcriptomic alterations observed in clinical studies of clear cell renal cell carcinoma (ccRCC), providing novel insights on the functional significance and therapeutic vulnerabilities of recurrent aneuploidies in human diseases.

## RESULTS

### Centromeric dCas9 recruitment induces efficient chromosome-specific mis-segregation and aneuploidy

The tumor-initiating event in ccRCC involves chromosome 3 (Chr3) aneuploidy^21^. To identify and isolate this rare Chr3 aneuploid population from tissue-derived normal renal epithelial cells, we adapted CRISPR LiveFISH^22^, a method that fluorescently labels repetitive genomic regions in live cells using dCas9 and fluorophore-tagged guide RNA (gRNA), for live-cell chromosome enumeration. While we successfully labelled Chr3 with dCas9 and a gRNA targeting Chr3-specific centromeric αSat HOR repetitive sequence, this approach also unexpectedly induced efficient Chr3-specific mis-segregation and micronucleus formation (Figure 1A). As over 70 binding sites of this gRNA are present across the Chr3 CENPA-bound region (Figure S1A), we hypothesized that dCas9-induced disruption at the kinetochore is responsible for this mis-segregation phenotype. Therefore, to exploit this finding for targeted aneuploidy engineering, we designed chromosome-specific gRNAs for all 24 human chromosomes, targeting their corresponding repetitive CENPA-bound regions as annotated by the T2T consortium^16–18^ (Figure 1B and S1A; Table S1).

**Figure 1.**
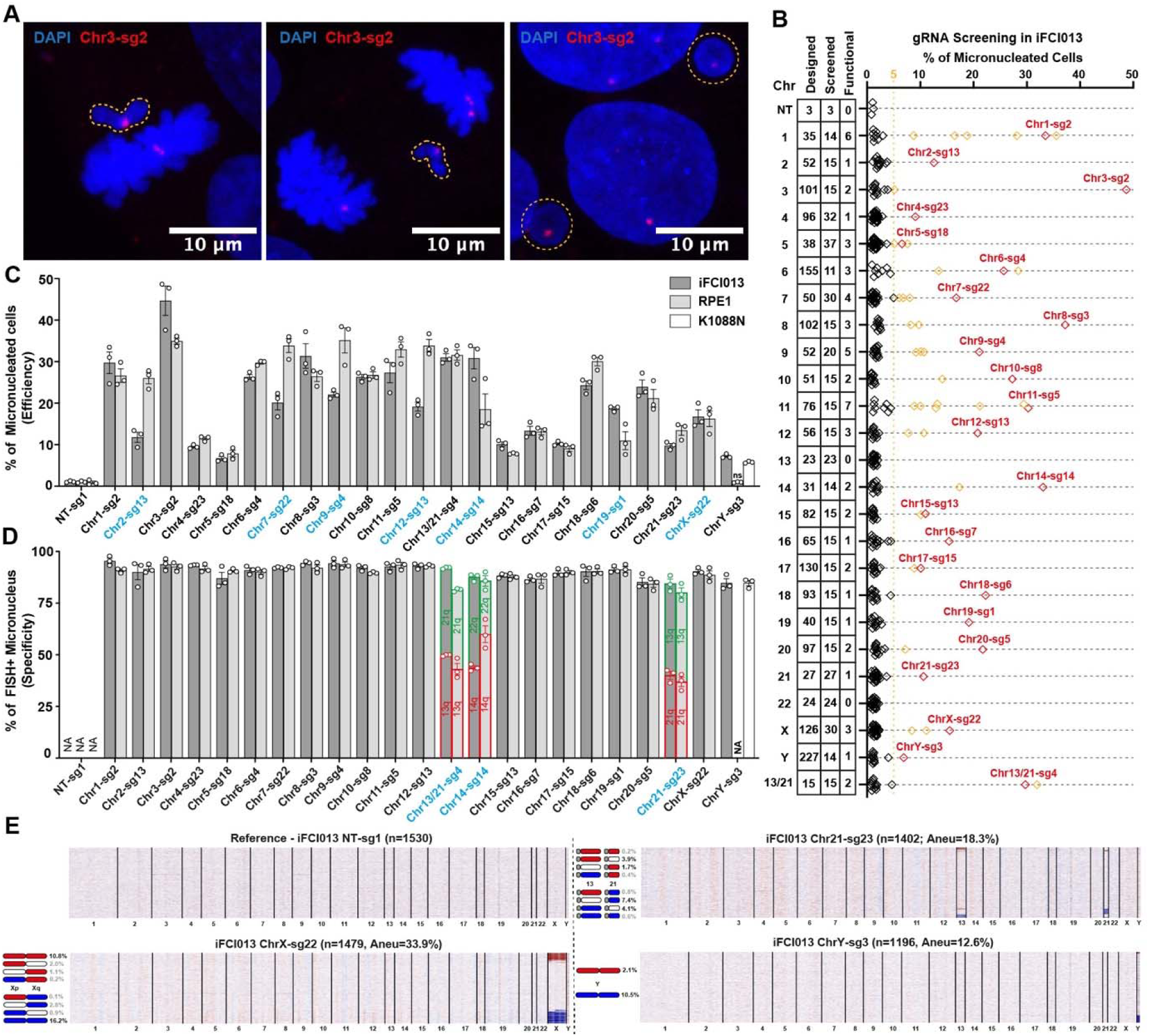
Centromeric dCas9 recruitment induces efficient chromosome-specific mis-segregation and aneuploidy. (A) Representative images showing Chr3-specific mitotic mis-segregation and micronucleus formation induced by dCas9 recruitment to Chr3-specific centromeric αSat HOR sequences with Atto550-labelled Chr3-sg2 gRNA in iFCI013 cells. Scale bars,10 μm. (B) Dot plot showing percentages of micronucleated cells induced by dCas9 with chromosome-specific gRNAs in iFCI013 cells at 48 h (Table S1). Rows represent chromosomes with the numbers of designed, screened, and functional gRNAs annotated. Diamonds represent gRNAs, and the orange line indicates a 5% threshold above which a gRNA is deemed functional. Functional gRNAs selected for validation are shown in red with gRNA ID labelled, and other functional gRNAs are shown in orange. (C) Bar plot showing percentage of micronucleated cells with the selected functional gRNAs in the indicated cell lines at 48 h. Functional gRNAs with model-specific efficiency differences are marked in blue. Error bars, mean ± SD (n=3 using different batches of gRNAs, >300 nuclei analyzed, one-way ANOVA followed by Dunnett’s post hoc multiple comparison test compared to the corresponding NT-sg1, all adjust p values < 0.01 except ChrY-sg3 in RPE1). (D) Bar plot showing percentage of chromosome-specific FISH positive (FISH+) micronucleus with the selected functional gRNAs in the indicated cell lines at 48 h. Functional gRNAs that target two chromosomes due to off-target effects or bi-chromosome-specific design are marked in blue, and their corresponding bars are split by target chromosomes shown in red and green. Error bars, mean ± SD (n=3, >100 micronuclei analyzed). (E) Heatmaps showing single-cell chromosome copy number profiles inferred from scRNA-seq data in iFCI013 cells after CRISPRt treatments. Each row represents an individual cell; columns correspond to chromosomes, scaled by the number of expressed genes. Copy number status for the target chromosome is shown in blue (loss), red (gain) or white (neutral). The gRNA used, the number of single cells analyzed by scRNA-seq (n), the percentage of all cells with aneuploidy of the targeted chromosome(s) (Aneu), and the percentage of cells with the indicated specific aneuploidy status are shown.

We tested 454 gRNAs for their ability to induce micronucleus formation in induced pluripotent stem cell (iPSC) line iFCI013 (Figure S1B). We identified 56 functional gRNAs (defined as inducing ≥5% micronucleated cells in 48 hours) for all chromosomes except Chr13 and Chr22 (Figure 1B and S2A). We validated the chromosome identity of the micronuclei induced by functional gRNAs using fluorescence *in situ* hybridization (FISH). While most functional gRNAs demonstrated on-target specificity, three gRNAs exhibited significant off-target effects: two Chr14 gRNAs (Chr14-sg11 and sg14) induced micronuclei for both Chr14 and Chr22, and one Chr21 gRNA (Chr21-sg23) induced micronuclei for both Chr21 and Chr13 (Figure S2A). Chr13/21 and Chr14/22 are two pairs of acrocentric chromosomes sharing high sequence homology within CENPA-bound regions^17^, and the observed off-target activities are likely driven by the identified off-target binding sites of these three gRNAs (Figure S2B). Given this sequence homology, we further designed 15 bi-chromosome-specific Chr13/21 gRNAs targeting the shared CENPA-bound sequence of Chr13 and Chr21 (Figure S2A; Table S1). We identified two functional Chr13/21 gRNAs that bi-specifically induce micronucleus formation for both Chr13 and Chr21 with markedly improved efficiency compared to Chr21-sg23 (Figure 1B and S2A).

Next, we selected a panel of 23 efficient functional gRNAs covering all 24 human chromosomes and further validated their performance in *TP53* proficient RPE1 retinal pigment epithelial cells (or K1088N primary renal epithelial cells for ChrY-sg3). We confirmed chromosome-specific micronucleus induction across different models using this gRNA panel (Figure 1C, 1D, and S3); however, the efficiency of several gRNAs varied between models (Figure 1C; see Discussion). We referred to this approach as ‘CRISPR-Taiji’ (CRISPRt), inspired by the imperfect segregation depicted in the ‘Taiji’ symbol.

To confirm aneuploidy generation via CRISPRt-induced mis-segregation, we performed single-cell RNA-seq (scRNA-seq) in iFCI013 iPSC cells after applying CRISPRt to Chr13, 21, X and Y, whose aneuploidies are associated with various chromosomal disorders (Figure S4A and S4B). The inferred single-cell copy number profiles using scRNA-seq data confirmed the generation of both gain and loss aneuploidies for the target chromosomes, without disrupting the expression of key pluripotency genes (Figure 1E, S4C and S4D). Notably, although Chr21-sg23 bi-specifically targeted both Chr13 and Chr21 as expected, their mis-segregation appeared to occur independently, generating predominantly single aneuploidy event of either chromosome in individual cells (Figure 1E).

In summary, we demonstrate that targeting dCas9 to the centromeric CENPA-bound regions induces efficient chromosome-specific mis-segregation and aneuploidy generation, applicable to all 24 human chromosomes.

### CRISPRt generates a diverse spectrum of chromosomal alterations

Human chromosomes differ in their propensity to mis-segregate, in part due to their nuclear positioning^23^. While this bias contributes to recurrent aneuploidy patterns in cancer, selection for aneuploidy-specific functional alterations is considered the dominant driver in shaping aneuploidy landscapes. However, a rapid and systematic approach to characterize phenotypic alterations across panels of recurrent aneuploidies in diverse biological contexts remains lacking. To address this question, we aim to leverage the efficiency, specificity and scalability of CRISPRt to engineer all autosomal whole-chr and chr-arm aneuploidies in isogenic backgrounds for phenotypic profiling.

We targeted all 22 autosomes for aneuploidy engineering using CRISPRt in near-euploid RPE1 and euploid RPTEC cells (both *TP53*-proficient), followed by scRNA-seq at five days following CRISPRt engineering (Figure 2 and S5A; see Star Methods for time point rationale). Inferred copy number analysis confirmed efficient chromosome-specific aneuploidy generation, achieving maximal on-target aneuploid cell proportion of 84.1% (median 56.4%) in RPE1 cells and 65.1% (median 38.7%) in RPTEC cells (Figure 2 and S5B). Across all targeted autosomes, micronucleated cell frequency quantified at 48 hours showed a robust linear correlation with the final aneuploid cell proportion (Figure S5C). Moreover, allele-specific expression analysis showed largely balanced representation of the remaining allele after chromosome loss in both RPE1 and RPTEC cells (Figure S5D and S5E), indicating limited haplotype targeting bias with these gRNAs.

**Figure 2.**
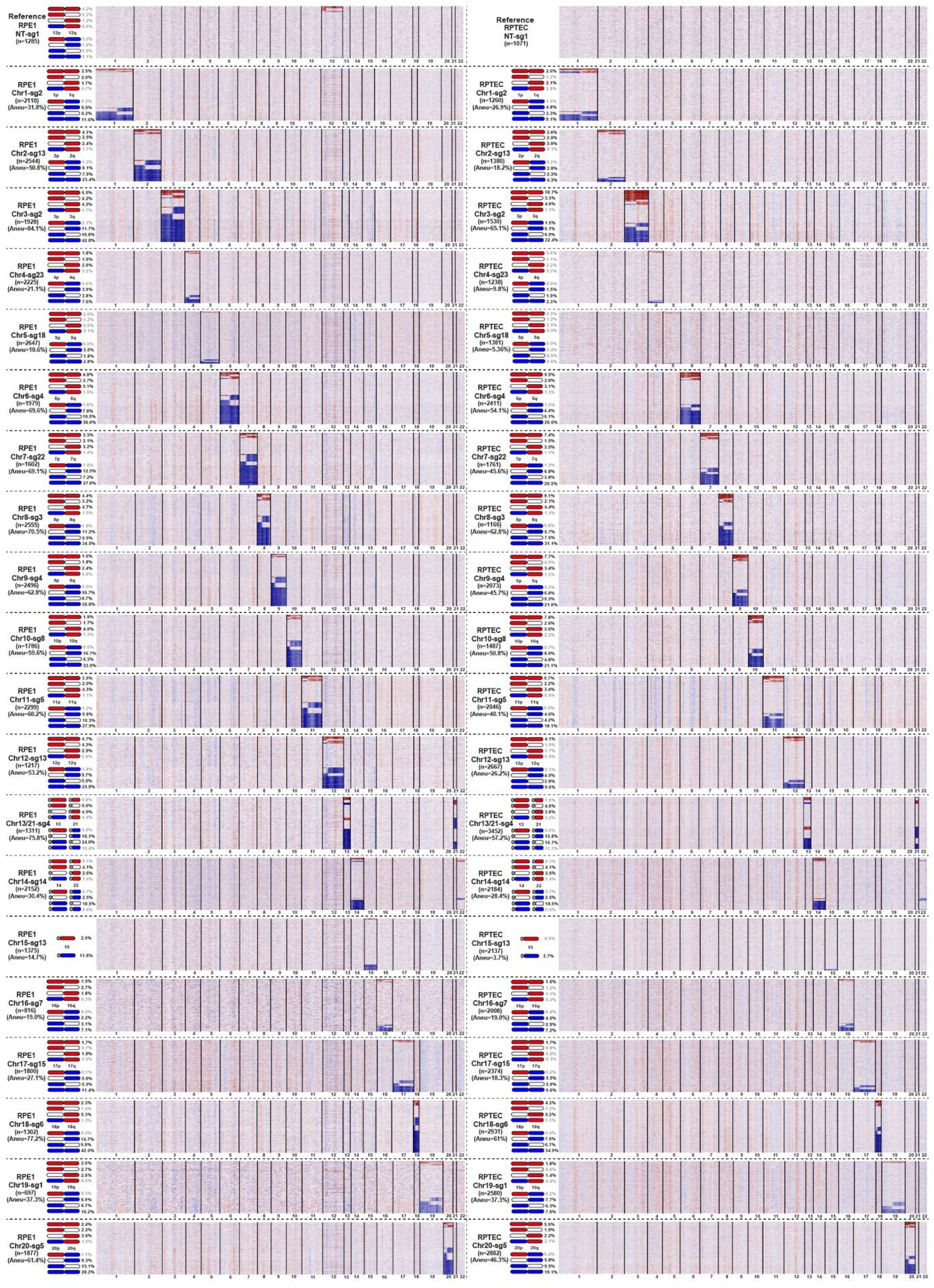
Pan-autosome CRISPRt aneuploidy engineering in RPE1 and RPTEC cells using CRISPRt. Heatmaps showing single-cell chromosome copy number profiles inferred from scRNA-seq data in RPE1 (left) and RPTEC (right) cells after CRISPRt treatments. Each row represents an individual cell; columns correspond to chromosomes, scaled by the number of expressed genes. Copy number status for the target chromosome is shown in blue (loss), red (gain) or white (neutral). The gRNA used, the number of single cells analyzed by scRNA-seq (n), the percentage of all cells with aneuploidy of the targeted chromosome(s) (Aneu), and the percentage of cells with the indicated specific aneuploidy status are shown.

Next, we analysed the CRISPRt-induced aneuploidy types across all targeted autosomes. Chromosomal losses were consistently more than gains, with RPE1 cells exhibiting a particularly pronounced bias toward losses (Figure S5F). Additionally, while whole-chr aneuploidies were predominant (>50% of events) as the expected mis-segregation outcome, chr-arm aneuploidies with putative centromeric breakpoints also appeared frequently (Figure S5G). We validated these aneuploidy types and confirmed centromeric breakage of chr-arm aneuploidies following Chr3 CRISPRt by chromosome painting in RPTEC-shP53 cells with stable *TP53* knockdown (Figure 3A and S6A; see Star Methods for cell line choice). Additionally, we observed that CRISPRt also induced frequent chromosome-specific shattering (chromothripsis) (Figure S6A and S6B), which underlies extensive genomic rearrangements and extrachromosomal DNA formation^24^.

**Figure 3.**
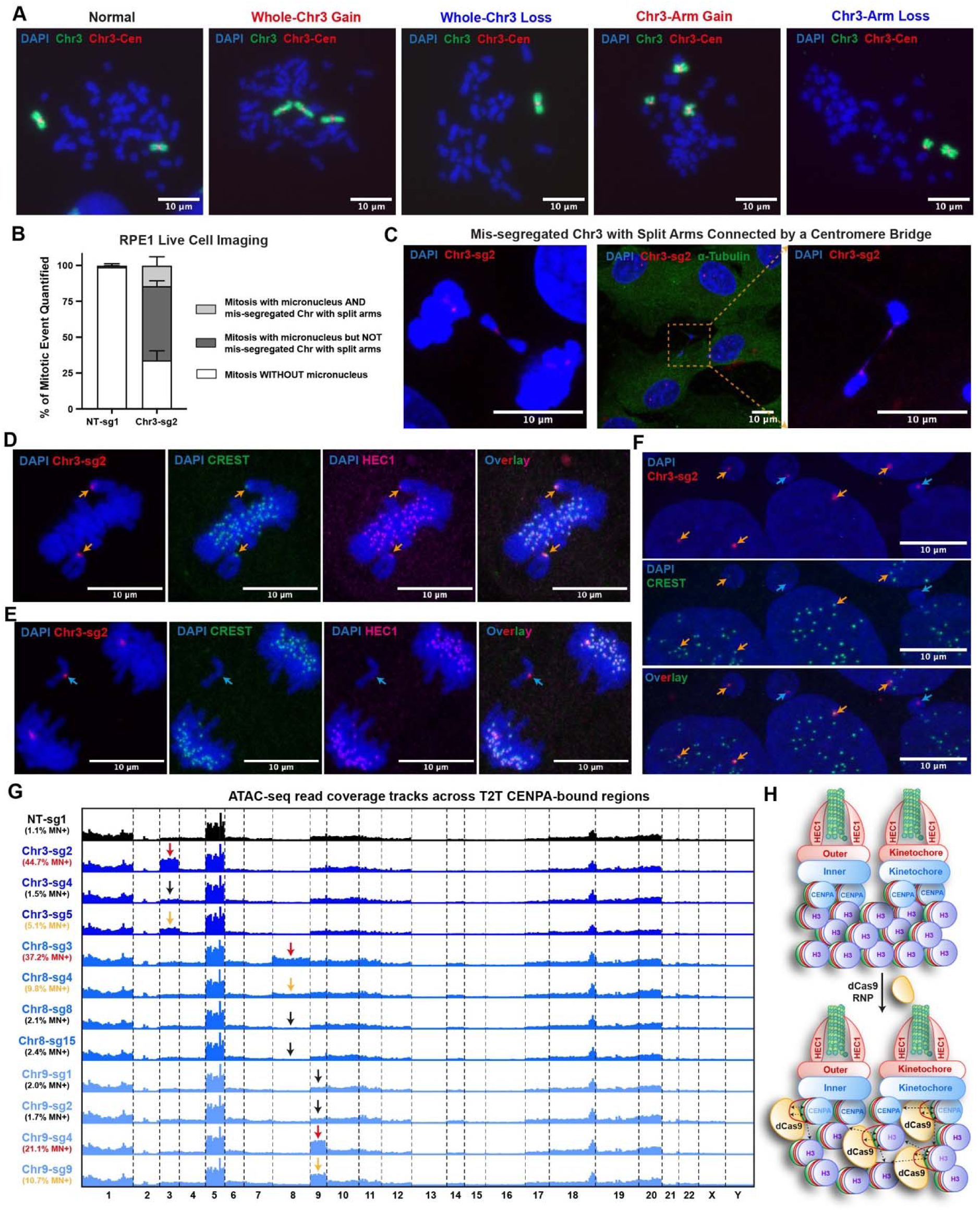
CRISPRt generates a diverse spectrum of chromosomal alterations via centromeric chromatin relaxation. (A) Representative images showing metaphase spreads with the indicated types of Chr3 aneuploid events 7 days after CRISPRt with Chr3-sg2 gRNA in RPTEC-shP53 cells. Chr3 arms and Chr3 centromere are shown in green and red, respectively. Scale bars,10 μm. (B) Bar plot showing percentage of mitotic events with the classified outcomes by time-lapse live-cell imaging (Video S1, see Star Methods) after CRISPRt with the indicated gRNAs in RPE1 cells. Error bars, mean ± SD (n=3, >200 mitotic events analyzed) (C) Representative images showing the mis-segregated Chr3 with split chromosome arms connected by the centromere bridge 24 h after CRISPRt with Atto550-labelled Chr3-sg2 gRNA in RPE1 cells. Chr3-sg2 (red) marks Chr3 centromere, and α-Tubulin (green) marks the cytoplasm. Scale bars,10 μm. (D, E and F) Representative images showing kinetochore components on Chr3 centromere in metaphase (D), anaphase (E), and interphase (F) 24 h after Chr3 CRISPRt with Atto550-labelled Chr3-sg2 gRNA in iFCI013 cells. Chr3-sg2 (red) marks Chr3 centromere, CREST antibody (green, stains for CENPA/B/C) marks the inner kinetochore, and HEC1 antibody (magenta) marks the outer kinetochore. Scale bars,10 μm. (G) Read coverage tracks showing representative normalized ATAC-seq signal across the CENPA-bound regions 8 h after CRISPRt with the indicated gRNAs in iFCI013 cells (n=2). Arrows indicate the targeted chromosomes. Micronucleus induction efficiencies of the gRNAs are shown, with colors reflecting efficiency. (H) Schematics showing dCas9 recruitment disrupts the centromeric chromatin organization at the CENPA-bound kinetochore region.

The formation of chr-arm aneuploidies requires centromeric DNA double-strand breaks, which dCas9 cannot directly generate. To understand the origin of CRISPRt-induced chr-arm aneuploidies, we performed live-cell imaging to analyse the fate of mis-segregated chromosomes. Following Chr3 CRISPRt, mis-segregation and micronucleus formation were observed in approximately two-thirds of mitotic events (Figure 3B). Notably, in over 20% of these abnormal mitoses, we observed mis-segregated chromosomes with split arms tethered by a centromeric DNA bridge spanning the two daughter cells (Figure 3B and 3C; Video S1). These centromeric bridges are susceptible to break under actomyosin contractile forces after cytokinesis^25^, thus leading to the generation of chr-arm aneuploidies. Therefore, CRISPRt-induced chr-arm aneuploidies likely originate from a subset of mis-segregated chromosomes exhibiting split-arm phenotypes.

In summary, we demonstrate that CRISPRt enables efficient engineering of both whole-chr and chr-arm aneuploidies across human autosomes.

### dCas9-induced centromeric chromatin relaxation underlies CRISPRt-induced chromosome mis-segregation

The generation of both whole-chr and chr-arm aneuploidies by CRISPRt prompted us to further investigate the mechanism that underlies CRISPRt-induced chromosome mis-segregation. We first explored potential kinetochore defects following CRISPRt treatment. In metaphase cells, Chr3 frequently showed misalignment at the metaphase plate, however, both the inner (CENPA/B/C indicated by CREST antibody staining) and outer (HEC1) kinetochore components were present (Figure 3D and S6C). More surprisingly, following anaphase onset, nearly all lagging Chr3 showed absence of both the inner and outer kinetochore components at their centromeres (Figure 3E and S6C). Consistently, following mitotic exit, 75% of Chr3 centromeres in interphase micronuclei remained devoid of inner kinetochore components (Figure 3F and S6C).

Notably, the same spectrum of mitotic defects has been observed following Polo-like kinase 1 (*PLK1*) inhibition, including defective maintenance of metaphase chromosome alignment, loss of the inner kinetochore, chromosome mis-segregations, and abnormal chromosomes with collapsed centromere and split arms^26,27^. Mechanistically, loss of *PLK1* function leads to aberrant centromeric recruitment of DNA helicases, driving centromeric DNA unwinding and chromatin relaxation^26,27^. This reduces structural rigidity of centromeric kinetochore chromatin, leading to the observed mitotic defects and chromosome mis-segregation under microtubule pulling forces^26,27^. Remarkably, dCas9 has helicase-like DNA unwinding activity and has been previously reported to induce chromatin relaxation at non-centromeric regions^28^. These together suggest that CRISPRt likely functions through centromeric chromatin disruption as well.

To test this hypothesis, we performed ATAC-seq to measure chromatin accessibility after CRISPRt using gRNAs with different micronucleus induction efficiencies. Indeed, all functional gRNAs, but not the non-functional ones, specifically induced accessibility increase at the targeted CENPA-bound centromeric regions, with non-centromeric regions unaffected (Figure 3G and S6D). Notably, functional gRNAs with higher micronucleus induction efficiency were associated with higher accessibility increase (Figure 3G and S6D). To elucidate whether dCas9 recruitment efficiency determines accessibility changes, we evaluated recruitment efficiency by quantifying the number and intensity of centromeric dCas9 foci recruited using the same set of gRNAs. Indeed, stronger dCas9 recruitment was associated with higher accessibility increase among the functional gRNAs (Figure S6D-F). Interestingly, non-functional gRNAs Chr8-sg15 and Chr9-sg1 exhibited comparable centromeric dCas9 recruitment but induced no accessibility increase (Figure S6D-F). This suggests that dCas9 recruitment is not the only determinant of chromatin relaxation and mis-segregation efficiency, while additional factors, including dCas9 binding orientation, may also be important.

Overall, our data demonstrate that dCas9-induced centromeric chromatin relaxation is the underlying mechanism of CRISPRt-induced mis-segregation and aneuploidy generation (Figure 3H).

### Comprehensive transcriptomic analysis of whole-chr aneuploidies reveals functionally significant alterations

The scRNA-seq data from CRISPRt-engineered cells provides not only the inferred aneuploidy status but also the paired transcriptomic profiles of individual cell. This enables rapid and systemic mapping of genotype-phenotype correlation across all engineered aneuploidies without the need to establish individual aneuploid clonal populations, analogous to the Perturb-seq strategy used to map the phenotypic consequences of gene-level perturbations at genome scale^29^.

We first focused on the transcriptomic alterations associated with individual engineered whole-chr aneuploidies. We excluded aneuploid populations representing <1.5% of total cells in the corresponding samples, resulting in 82 whole-chr loss or gain populations from RPE1 and RPTEC cells for downstream analysis. The primary transcriptomic consequence of aneuploidy is gene dosage alterations across the affected chromosome, which further induce secondary genome-wide effects^30^. To evaluate these effects, we performed gene set enrichment analysis (GSEA) using hallmark and KEGG gene sets, comparing each aneuploid population to its corresponding copy-neutral counterpart, with or without excluding genes on the aneuploid chromosome (Figure 4A and S7A). We found excluding these genes had minimal impact on pathway-level alterations (Figure 4A and S7A), suggesting that the observed changes primarily result from secondary genome-wide effects of aneuploidies, which we further explored in the subsequent analyses.

**Figure 4.**
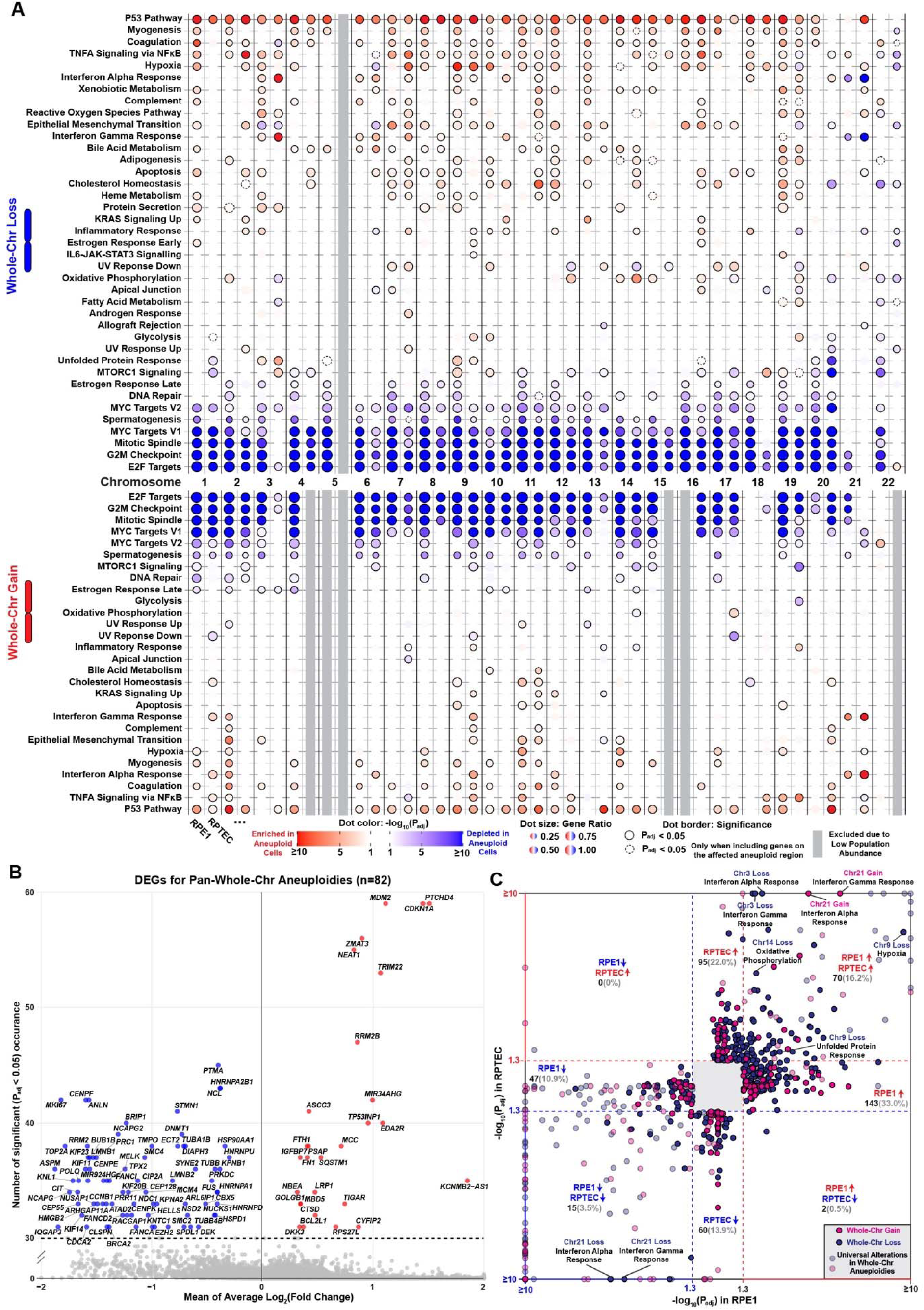
Transcriptomic analyses reveal universal and aneuploidy-specific transcriptomic responses to whole-chr aneuploidies. (A) Dot plot showing significantly enriched or depleted hallmark gene sets by gene set enrichment analysis (GSEA) in RPE1 and RPTEC cells with the indicated whole-chromosome aneuploidy, compared to the corresponding copy-neutral population. The main analyses exclude genes located within the corresponding aneuploid region to highlight the secondary genome-wide effects. The columns represent cell type and chromosomes. The rows represent significantly altered hallmark gene sets, ordered by average enrichment scores. The color gradient represents –log_10_(P_adj_) values with red and blue corresponding to positive and negative normalized enrichment scores, respectively. Dots with a solid border highlight significant changes (P_adj_ <0.05). Dots with a dashed border highlight additional significant changes when all the genes are included in the analyses for the global changes. (B) Volcano plot showing differentially expressed genes (DEGs) across the 82 whole-chromosome aneuploid populations shown in Figure 4A. DEG lists are first generated separately for each aneuploid population and then aggregated to compute the mean Log₂(Fold Change) and the number of significant occurrences (P_adj_ <0.05) across all 82 populations. DEGs with consistent significant occurrence (≥30) are labelled with HGNC symbols. (C) Scatter plot showing shared or cell line-specific hallmark gene set alterations in RPE1 and RPTEC cells with whole-chromosome aneuploidy. Each dot represents a significantly altered gene set for a specific aneuploid population, with highlighted gene sets labelled by name and associated aneuploid event. The x– and y-axes show the corresponding –log_10_(P_adj_) values in RPE1 and RPTEC cells, respectively, with color indicating the direction of enrichment. Transparent dots denote universal alterations in whole-chr aneuploidies including “P53 pathway”, “E2F targets”, “G2M checkpoint”, “Mitotic spindle”, “MYC targets V1/2”, and “Spermatogenesis”. The number and percentage of gene set alterations, excluding the universal alterations, with the indicated alterations in RPE1 and RPTEC cells are shown.

We first characterised alterations consistently observed across engineered aneuploid populations, representing the universal transcriptomic response to whole-chr aneuploidy. We found that activation of *TP53* pathway and downregulation of cell cycle-related gene sets (*E2F* targets, G2M checkpoint, mitotic spindle, *MYC* targets V1/V2, and spermatogenesis) comprise this universal response (Figure 4A and 4B), consistent with previously described aneuploidy-associated stress phenotype^31^. We noted that *TP53* pathway may also be activated by the prolonged mitotic duration following CRISPRt treatment^32^ (Figure S7B). Additionally, significant overlaps among the GSEA leading-edge genes of the cell cycle–related gene sets may also partially explain their concordant downregulation (Figure S8C). Previous studies reported consistent downregulation of ribosome biogenesis in established monosomic or polysomic clones^33,34^. However, this alteration was only observed with a limited subset of our engineered whole-chr aneuploidies, while upregulation of lysosome pathway was more universal, particularly for whole-chr losses (Figure S7A). These differences may reflect a dynamic adaptation to aneuploidy-induced proteotoxic stress^31,34^, as our experiment captured early responses following aneuploidy induction, whereas previous studies analysed long-term expanded aneuploid clones.

Besides these universal alterations, the other changes are aneuploidy-specific with variations across cell lines, likely driving distinct functional outcomes. Over 75% of these alterations are unique to either RPE1 or RPTEC cells (Figure 4C), highlighting cell lineage as a key determinant, possibly due to differences in baseline gene expression and regulatory networks (Figure S7D). These lineage-specific alterations may underlie the distinct aneuploidy patterns observed across diverse cancer types. In contrast, the remaining 23% of aneuploidy-specific alterations are consistent across both cell lines (Figure 4C), likely reflecting conserved functional changes across broader cellular contexts. Among these lineage-conserved aneuploidy-specific alterations, we observed significant upregulation of interferon responses associated with Chr3 and Chr6 losses, as well as Chr21 gain (Figure 4A and S8A-C). Notably, activated interferon response is a key cellular phenotype of Down syndrome (Chr21 gain), driven by increased dosage of interferon receptor genes located on Chr21^35^ (Figure S8D). Importantly, engineered Chr21 loss exhibited the opposite effect, with significant downregulation of interferon responses (Figure 4A, S8D and S8E).

As our approach robustly recapitulated well-characterized changes associated with Chr21 gain, we confidently proceeded to analyze additional observed aneuploidy-specific alterations and explore their potential disease relevance. We focused on Chr9 and Chr14 losses, which are strongly associated with the acquisition of metastatic competence in ccRCC^36^. The most significant alteration associated with engineered Chr9 loss was upregulation of hallmark hypoxia gene set (Figure 4A and S8F), which was further validated using a stringent hypoxia-inducible factor (HIF) target gene set^37^ (Figure S8G). This highlights a strong correlation between Chr9 loss and HIF signaling activation, despite the underlying mechanism remaining unclear. However, this association was absent in our multi-omics profiling analyses of ccRCC tumor samples^38^, likely masked by constitutive HIF signaling activation due to near-universal *VHL* loss in ccRCC^39^. Instead, upregulated unfolded protein response (UPR) was the most significant alteration observed in ccRCC tumor regions with Chr9 loss^38^. Notably, engineered Chr9 loss in both cell lines recapitultated significant UPR upregulation (Figure 4A and S8H), suggesting a potential lineage-conserved function of Chr9 loss in buffering proteotoxic stress. Next, among all engineered aneuploidies, only Chr14 loss induced lineage-conserved upregulation of oxidative phosphorylation gene set (Figure 4A and S8I). This feature was recently reported in ccRCC tumors with Chr14 loss^40^. More importantly, increased oxidative metabolism is observed in human ccRCC metastatic sites and directly drives metastasis in syngeneic mouse model of ccRCC^41^, highlighting a potential mechanism for Chr14 loss in promoting metastasis.

In summary, we systematically characterized the transcriptomic alterations of individual whole-chr aneuploidies in isogenic backgrounds, providing a framework to identify potential disease-relevant alterations driven by specific aneuploidies.

### Dissecting aneuploidy-specific alterations into chr-arm and gene-level effects

Both whole-chromosome and arm-level aneuploidies are prevalent in cancer, but individual chromosomes often show recurrent aneuploidy patterns that affect the p-arm, q-arm, or the entire chromosome differently, suggesting distinct functional consequences. To explore this, we compared transcriptomic responses between whole-chr aneuploidies and their corresponding p-arm and q-arm aneuploidies.

After removing chr-arm aneuploid populations with low engineering efficiency, we performed GSEA using hallmark gene sets, excluding genes located on the corresponding aneuploid arm (Figure 5A and S9A). First, the universal aneuploidy-associated stress response was generally conserved across chr-arm aneuploidies. Then, we compared aneuploidy-specific alterations between whole-chr and corresponding chr-arm aneuploidies to delineate their shared and distinct transcriptomic effects. Across significant alterations associated with whole-chr aneuploidies, over 60% showed concordant changes in the corresponding chr-arm aneuploidies, with more than half attributed to either the p-arm or q-arm specifically (Figure 5B and S9B). This suggests that individual chromosome arms have both common and distinct contributions that substantially shape the alterations for the corresponding whole-chr aneuploidy. For example, major alterations associated with Chr6 loss, including upregulation of interferon responses, appear to be driven specifically by Chr6p loss, but not Chr6q loss (Figure 5A). Interestingly, Chr6q loss is prevalent across cancer types^42^, but losses of Chr6 or Chr6p are rare, despite frequent focal deletion of the human leukocyte antigen (HLA) locus on Chr6p^43^. Our observations suggest negative selection against losses of Chr6 or Chr6p, potentially mediated by specific observed alterations associated with Chr6p loss. Additionally, unique alterations associated with either whole-chr or chr-arm aneuploidies were also common (Figure 5B), likely reflecting some synergistic or compensatory interactions between the p-arm and q-arm.

**Figure 5.**
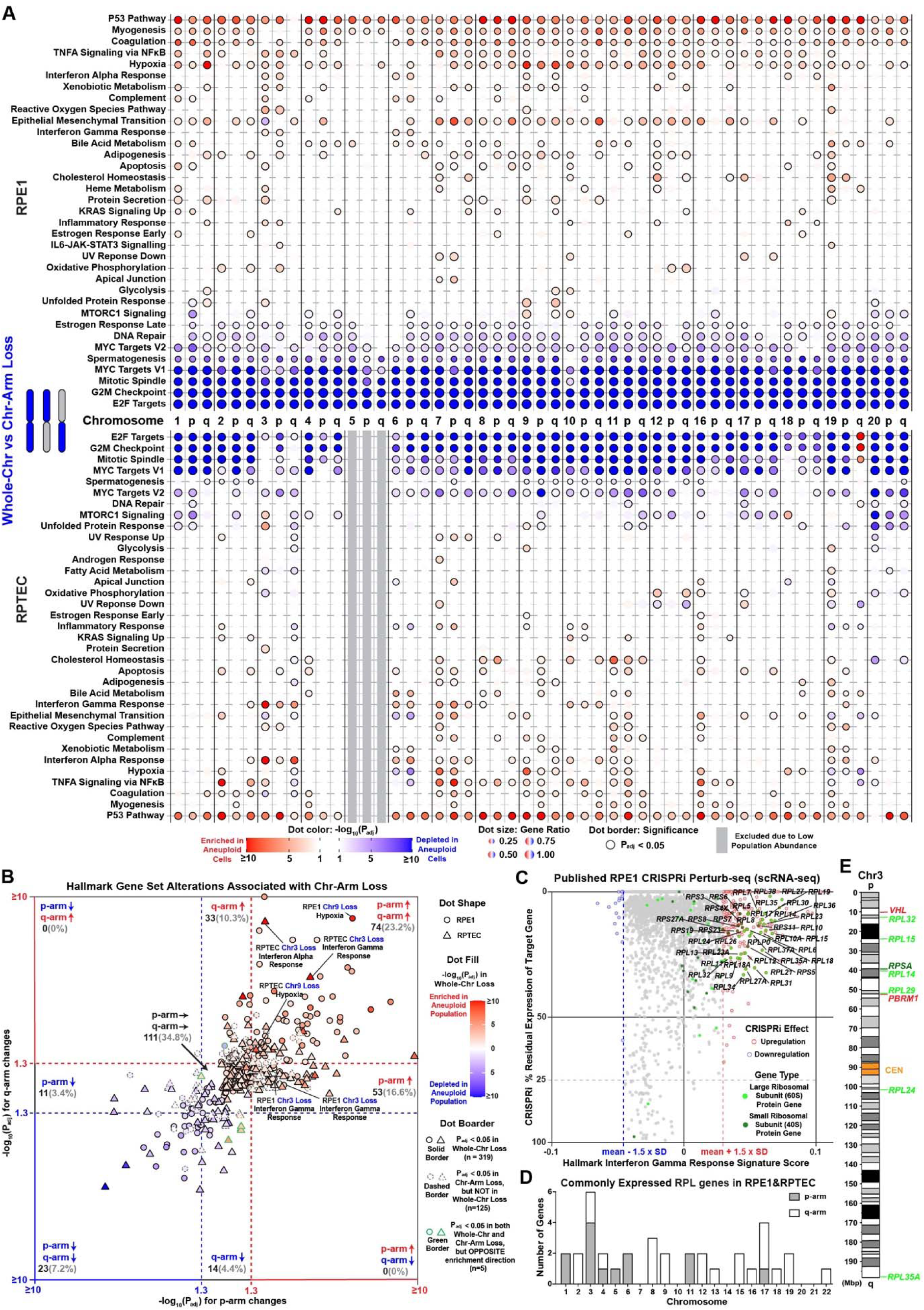
Chr-arm and gene-level changes likely underlie specific alterations associated with whole-chr aneuploidies. (A) Dot plot showing significantly enriched or depleted hallmark gene sets by GSEA in RPE1 and RPTEC cells with indicated whole-chr and chr-arm losses, compared to the corresponding copy-neutral population. The analyses exclude genes located within the corresponding aneuploid region to highlight secondary genome-wide effects. The columns represent the indicated whole-chr or chr-arm loss. The color gradient represents –log_10_(P_adj_) values with red and blue corresponding to positive and negative normalized enrichment scores, respectively. Dots with a solid border highlight significant changes (P_adj_ <0.05). (B) Scatter plot showing the hallmark gene set alterations associated with the corresponding chr-arm losses in RPE1 (circles) and RPTEC (triangles) cells, excluding the universal alterations. Each dot represents an altered gene set associated with whole-chr or chr-arm loss of a specific chromosome in RPE1 or RPTEC cells, with highlighted gene sets labelled by name, cell type and chromosome. The x– and y-axes show the –log_10_(P_adj_) values for the corresponding p-arm and q-arm losses, respectively, with color indicating the direction of enrichment. Dot fill color shows the –log_10_(P_adj_) values and the direction of enrichment observed in the corresponding whole-chr loss. Border types represent different combinations of significance and enrichment directions in the corresponding whole-chr and chr-arm losses. The number and percentage of significant gene set alterations in the whole-chr loss with the indicated changes in the corresponding chr-arm losses are shown. (C) Scatter plot showing the correlation between CRISPRi-induced gene perturbations and hallmark interferon gamma response signature scores with the published RPE1 essential-wide Perturb-seq dataset^29^. The x– and y-axes show the signature score and the residual expression of the target gene after CRISPRi knockdown. Ribosomal large subunit protein (RPL) and small subunit protein (RPS) genes with a significant effect on interferon gamma response are labelled. (D) Histogram showing the distribution of consistently expressed RPL genes in RPE1 and RPTEC cells across chromosome arms. (E) Schematics showing the genomic positions of RPL and RPS genes as well as two tumor suppressor genes “*VHL*” and “*PBRM1*” on Chr3. Orange boxes represent Chr3 centromere.

Following comprehensive mapping of aneuploidy-specific alterations at chr-arm level, we further attempted to identify their underlying gene-level determinants for potential therapeutic implications. To do this, we leveraged published datasets of CRISPR interference (CRISPRi) Perturb-seq, which maps gene-level phenotypic consequences following knockdown of individual genes at a genome-wide scale. Given the diverse cell types used in published Perturb-seq datasets, we focused on the lineage-conserved upregulation of interferon responses associated with Chr3p loss (Figure 5A), which was also previously observed in lung squamous cell carcinoma^7^, indicating a broadly conserved mechanism. Using three published Perturb-seq datasets, we identified genes whose knockdown led to increased or decreased interferon gamma response^29,44^ (Figure 5C, S9C and S9D), however, these genes showed no clear distribution bias towards specific chromosome arms (Figure S9F). Further gene ontology analysis revealed significant enrichment of cytosolic large (RPL) and small (RPS) ribosomal subunit genes among those whose knockdown activated interferon gamma response (Figure 5C, S9C and S9D). This is consistent with activated interferon signaling observed in early erythroid progenitors from patients with Diamond-Blackfan anemia carrying germline mutations in RPL or RPS genes^45^. Interestingly, among all chromosome arms, Chr3p encompasses the most constitutively expressed RPL genes, along with an additional RPS gene (Figure 5D, 5E and S9G). Therefore, the concordant, modest dosage reduction of multiple ribosomal genes on Chr3p (Figure S9H) likely exerts synergistic effects that drive pronounced upregulation of interferon responses.

Overall, we reveal both distinct and shared transcriptomic responses between whole-chr and corresponding chr-arm aneuploidies and demonstrate the identification of candidate driver genes using gene-level perturbation data.

### Deciphering early tumorigenic programs induced by the ccRCC-initiating aneuploidy event

Recent studies have identified expanded aneuploid normal epithelial cells lacking oncogenic mutations across various tissues^3–6^. Remarkably, some of these aneuploidies show similar recurrent patterns observed in corresponding cancer types^4,5^ suggesting that pre-existing aneuploidies in normal cells may contribute to tumorigenesis. A well-established example is Chr21 trisomy, which predisposes children with Down syndrome to acute leukaemia^46^. However, the potential tumorigenic roles of many other recurrent aneuploidies identified in normal cells remain largely unknown. Therefore, we sought to leverage CRISPRt’s simplicity and efficiency in targeted aneuploidy engineering of *TP53*-proficient normal cells to explore this question.

Here, we focused on Chr3p loss in normal renal proximal tubular epithelial cells (PTECs), the first tumor-initiating genetic alteration in nearly all ccRCC cases^21^ (Figure S10A). Chr3p loss in ccRCC minimally spans from p-arm telomere to *PBRM1* gene (Figure 5E) and can extend into Chr3q or the entireChr3^21,47^. For simplicity, we refer to these collectively as ‘Chr3(p) loss’ hereafter. Chr3(p) loss leads to heterozygous deletion of multiple tumor suppressor genes, including *VHL*, which encodes a master regulator of HIF signalling^48^ (Figure S10B). In ccRCC tumors, inactivation of the remaining wild-type (WT) *VHL* allele through mutation or methylation occurs near-universally following Chr3(p) loss, leading to *VHL* biallelic loss (*VHL* loss) and constitutive activation of oncogenic HIF signalling^49^ (Figure S10A and S10B). Together, these two genetic alterations are observed in over 90% of sporadic ccRCC cases^21^. Notably, *VHL* loss alone by biallelic knockout (KO) failed to initiate ccRCC in mouse models^50,51^, highlighting a crucial but unclear tumorigenic role of Chr3(p) loss. To faithfully recapitulate ccRCC tumorigenic steps, we engineered Chr3(p) and *VHL* loss in primary PTEC organoids K1088N, derived from a donor carrying a germline *VHL* mutation (Figure S10C and S10D). Chr3 CRISPRt (Chr3-Ct) in K1088N cells can generate two Chr3(p) loss populations, with or without concomitant *VHL* loss, depending on which Chr3 allele is lost (Figure S10D). This enabled simultaneous modelling of both key genotypes involved in ccRCC initiation (Figure S10A). Additionally, we performed *VHL* WT allele-specific KO (*VHL*-ASK) to model *VHL* loss alone, without Chr3(p) loss (Figure S10C-F).

We confirmed efficient generation of Chr3(p) loss (predominantly whole-Chr3 loss) at 7 days (D7) following Chr3-Ct using scRNA-seq (Figure 6A). In contrast, *VHL*-ASK generated almost no detectable Chr3 aneuploidies (Figure S10G). Despite Chr3(p) loss being the tumor-initiating event, cell proportion with Chr3(p) loss declined significantly after one additional passage at 11 days (D7+4) following Chr3-Ct (Figure 6A), which could reflect short-term fitness loss due to aneuploidy-associated stress response^31^. Therefore, to restore the Chr3(p) loss population for prolonged culture, we employed a phenotypic selection strategy by sorting for CA9-positive (CA9+) cells, a well-established membrane marker upregulated upon *VHL* loss (Figure S10D). Sorting for CA9+ cells from the *VHL*-ASK population confirmed effective enrichment of cells with a high HIF signature score, indicative of *VHL* loss (Figure S10H and S10I; See Methods). Similarly, sorting for CA9+ cells from the Chr3-Ct D7+4 population significantly enriched cells with Chr3(p) and *VHL* loss (Figure 6A, S10H and S10I). However, single-cell expansion of the enriched population still failed to yield any transformed clonal organoids that escape senescence (see Discussion).

**Figure 6.**
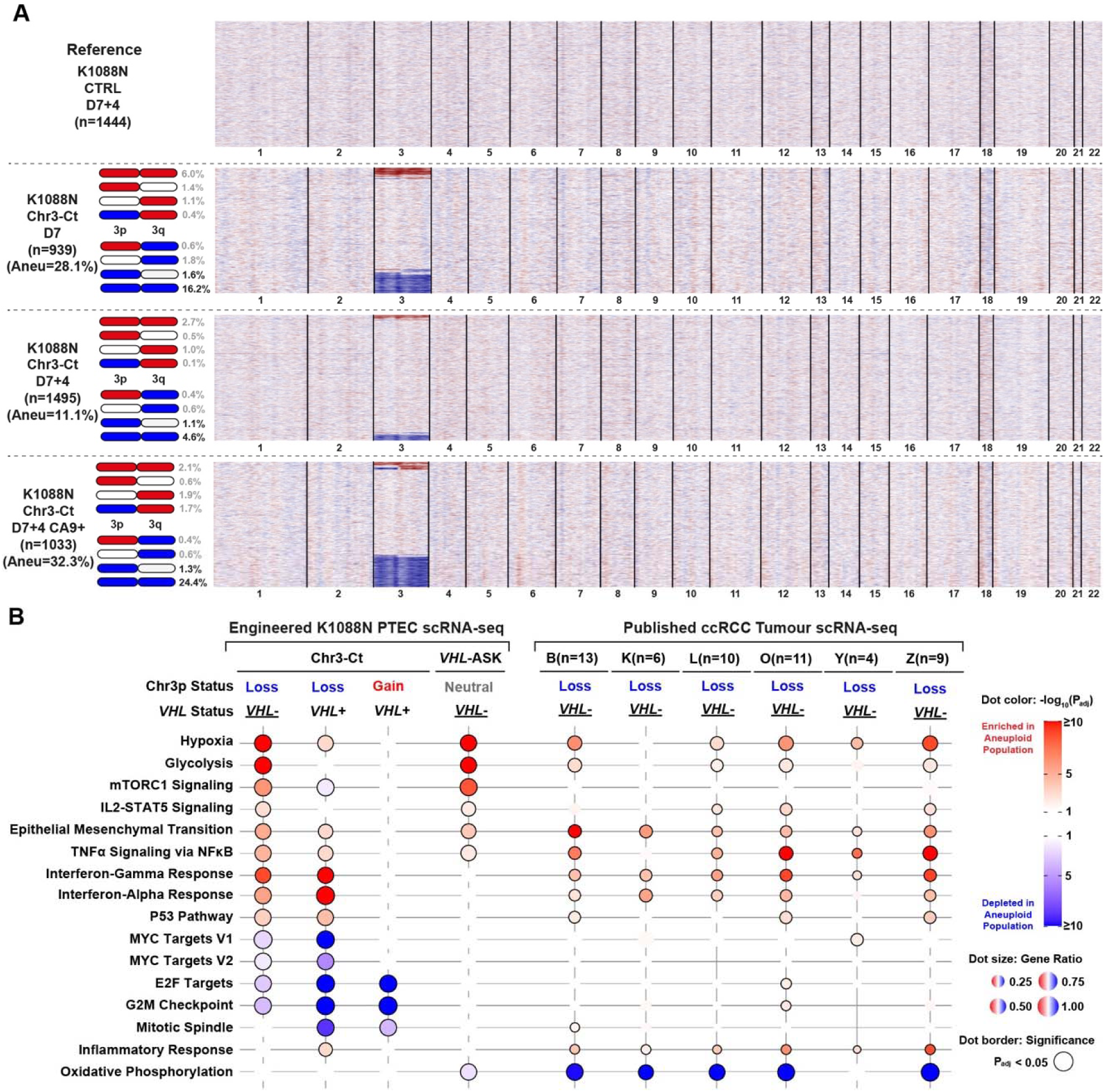
Tumor-initiating Chr3(p) loss drives upregulation of interferon responses as a consistent feature in ccRCC tumors. (A) Heatmaps showing single-cell chromosome copy number profiles inferred from scRNA-seq data of K1088N primary PTEC organoid cells with the indicated treatments. Each row represents an individual cell; columns correspond to chromosomes, scaled by the number of expressed genes. Copy number status for the target chromosome is shown in blue (loss), red (gain) or white (neutral). The total number of cells analyzed (n), the total percentage of cells with target aneuploid events (Aneu), and the percentage of cells with the indicated target aneuploidy status are shown. (B) Dot plot showing significantly enriched or depleted hallmark gene sets by GSEA in the engineered K1088N cells with the indicated genotypes (Figure S11A) and ccRCC cells from patient tumor samples, compared to the PTECs in the corresponding normal adjacent tissue. The abbreviations refer to the published ccRCC tumor scRNA-seq cohorts (B^52^, K^53^, L^54^, O^55^, Y^56^, and Z^57^), with the number of ccRCC tumors analyzed in each cohort (n) indicated. The analyses exclude genes located within the corresponding aneuploid region to highlight secondary genome-wide effects. The color gradient represents –log_10_(P_adj_) values with red and blue corresponding to positive and negative normalized enrichment scores, respectively. Dots with a solid border highlight significant changes (P_adj_ <0.05).

Nevertheless, we proceeded to analyze the transcriptomic consequences of Chr3(p) loss and *VHL* loss (Figure 6B and S11A). Notably, our engineered K1088N cells with both Chr3(p) and *VHL* loss recapitulated key transcriptomic features of ccRCC cells observed across six independent ccRCC tumor scRNA-seq studies^52–57^ (Figure 6B), demonstrating the fidelity of our model. We next analyzed our engineered K1088N cells with either Chr3(p) or *VHL* loss alone to deconvolve their distinct transcriptomic impacts on ccRCC initiation. Consistent with prior studies, *VHL* loss resulted in the dysregulation of hypoxia and glycolysis pathways, mTORC1 signaling, epithelial mesenchymal transition, and TNFα signalling^58–60^ (Figure 6B). Remarkably, Chr3(p) loss, but not *VHL* loss, resulted in the upregulation of interferon responses (Figure 6B). This alteration was consistently observed in ccRCC cells from tumors (Figure 6B), as well as in RPE1 and RPTEC cells with engineered Chr3(p) loss (Figure 5A and S11B). Interestingly, ccRCC is among the most immune-infiltrated solid tumors and the underlying mechanism remains unclear^61,62^. Our observation provides a plausible explanation that the near-universal Chr3(p) loss shapes this unique ccRCC immune phenotype via upregulated interferon responses. Moreover, immune cell influx and cytokine release have been shown to promote transformation of EGFR-mutant lung alveolar epithelial cells by inducing a progenitor-like cell state^63^, implying a potential contribution of the immune compartment in ccRCC initiation.

Furthermore, ccRCC tumors are highly immunogenic despite a low mutational burden and respond exceptionally well to immune checkpoint inhibition^64^. Recent work has demonstrated that *VHL* loss activates aberrant expression of endogenous retroviral elements, which may act as immunogenic tumor-associated antigens (TAAs) in ccRCC^38,65^. Interferon signaling is known to promote antigen presentation by upregulating antigen presentation machinery genes. Consistently, we observed significant upregulation of major histocompatibility complex (MHC) class I genes in both K1088N cells with engineered Chr3(p) loss and ccRCC tumor cells from six independent studies (Figure S11C and S11D). These together imply that the two near-universal initiating events of ccRCC may synergistically underlie its highly immunogenic phenotype.

Overall, we modelled the ccRCC-initiating Chr3(p) loss event in primary normal PTEC cells using CRISPRt and identified interferon response upregulation as a potential early tumourigenic program driven by Chr3(p) loss.

## DISCUSSION

We present CRISPRt, an approach to induce chromosome-specific mis-segregation and aneuploidy generation by targeting dCas9 to the core centromeric region. Despite its simplicity, CRISPRt achieves high efficiency and specificity, enables broad chromosome targetability, and is compatible across multiple model systems. It is also versatile in inducing various aneuploidy types, offering a generalizable solution for targeted aneuploidy engineering in diverse preclinical models. For a few chromosomes, we were unable to identify specific or highly efficient gRNAs despite exhaustively testing every design, which might be overcome by using other catalytically dead Cas variants with improved fidelity or broader PAM compatibility. Additionally, certain gRNAs showed inconsistent efficiencies between cell models, potentially due to sequence and location heterogeneities of the CENPA-bound regions^66,67^, warranting further investigations.

Mechanistically, relaxed centromeric chromatin underlies the centromere and kinetochore defects induced by CRISPRt. Notably, these defects, particularly loss of the inner kinetochore components, indicate the involvement of mechanical pulling forces from microtubules. Consistently, in *PLK1* inhibition studies, low concentrations of nocodazole or paclitaxel, which reduce microtubule attachment and its pulling forces, effectively rescued the same set of defects^26,27^. This represents a key difference between CRISPRt and KaryoCreate, as KaryoCreate was designed to induce mis-segregation by disrupting kinetochore-microtubule attachment^20^. However, only partial restoration of CRISPRt-induced inner kinetochore loss was observed (Figure 3F and S6C), likely causing the bias towards chromosome losses. Further modifications, such as fusing HJURP to dCas9, may improve kinetochore reestablishment for more efficient chromosome gain engineering^68^.

Using CRISPRt, we engineered and characterized individual aneuploidies across all 22 autosomes, revealing causal relationships between specific aneuploidies and disease-relevant transcriptomic alterations. For example, Chr9 loss activated unfolded protein response with significant upregulated *HSP90B1* (Figure S8H), a chaperone targetable by HSP90 inhibitors. Further comparative analysis revealed both shared and distinct alterations between whole-chr and corresponding chr-arm aneuploidies, which may underlie the selection of specific aneuploidy patterns. Moreover, while gene-level perturbation datasets can help identify candidate gene drivers of aneuploidy-induced alterations, precise experimental manipulation of multigene dosage for validation remains challenging. Nevertheless, emerging computational perturbation models may offer a powerful general solution for deciphering aneuploidy response and deconvoluting gene-level contributions directly from baseline transcriptomic profiles. Importantly, our work establishes a novel experimental framework for generating genome-wide aneuploidy-level perturbation datasets, providing crucial information for computational model training.

Lastly, it remains mostly unclear in various tissue contexts how specific aneuploidy in normal epithelial cells might contribute to tumorigenesis. Leveraging the broad model compatibility of CRISPRt, we demonstrate efficient engineering of Chr3(p) loss in tissue-derived normal epithelial cells and identified interferon response activation as a putative early tumorigenic program driven by Chr3(p) loss in the context of ccRCC. This implies an important role for immune components in promoting ccRCC tumorigenesis and may also account for the inability of biallelic *VHL* knockout to initiate ccRCC in mouse models^50,51^. Subsequent systematic identification of permissive factors that sustain PTECs with Chr3(p) loss is imperative to establish clonal Chr3(p) loss population for detailed functional characterization.

Overall, simply with dCas9, CRISPRt provides a practical and scalable approach for engineering and characterizing aneuploidies across diverse biological contexts.

### Limitations of the study

We leveraged inferred copy number profiles from scRNA-seq data to assign aneuploidy status. Although, we used the corresponding non-targeting control cells as the reference population to ensure more accurate copy number inference, this approach still lacks a definitive copy number ground truth and remain susceptible to false calls. Consequently, we excluded some aneuploid populations with low representation from further analysis. Recent technological advances, such as WellDR-seq^69^, which enables scalable single-cell whole-genome and transcriptome co-profiling, represent a better method for establishing a more accurate and comprehensive aneuploidy genotype–phenotype landscape when paired with CRISPRt engineering.

## FIGURE TITLES AND LEGENDS

**Figure S1.**
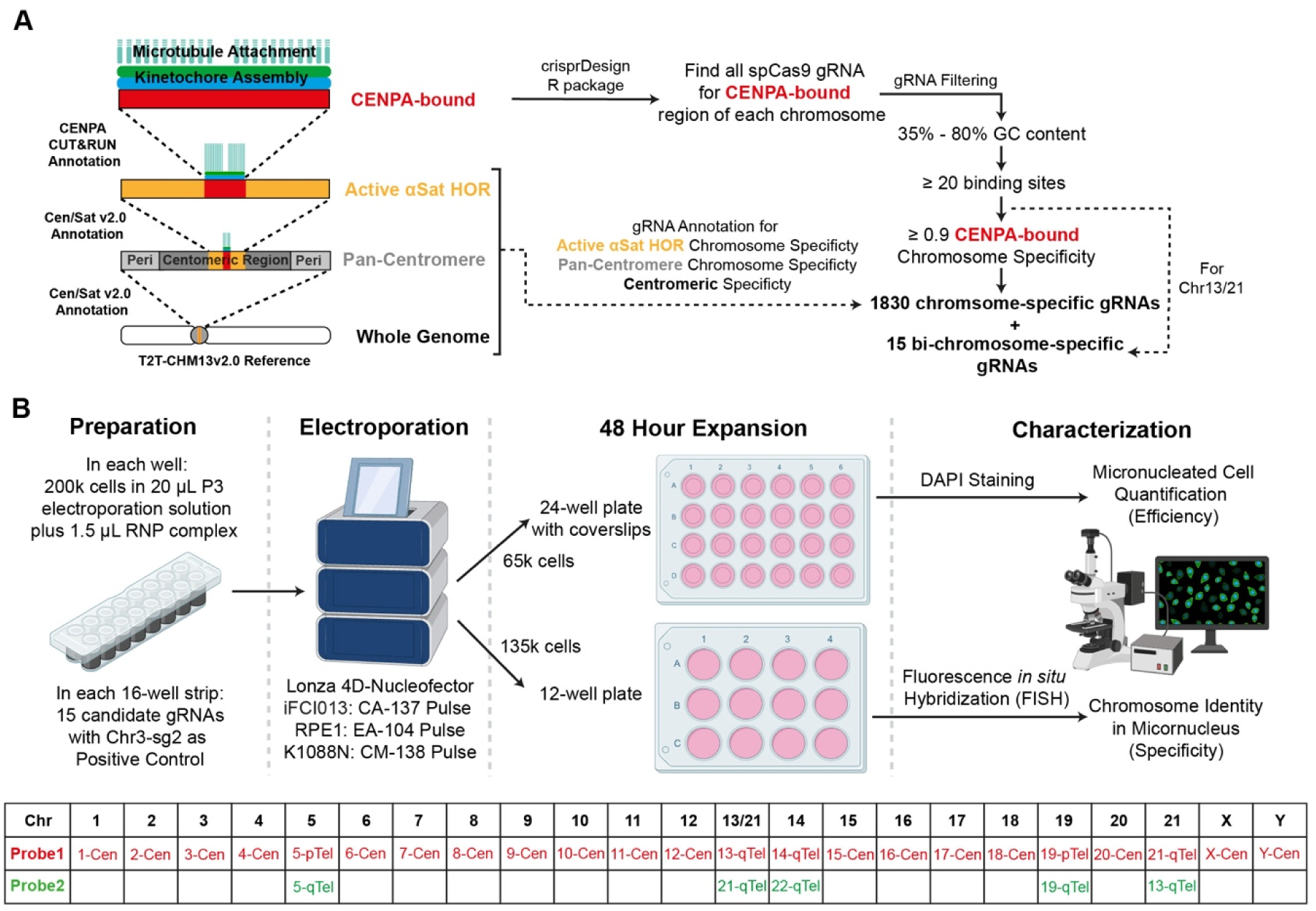
Design and screening of chromosome-specific gRNAs targeting CENPA-bound region, related to Figure 1. (A) Schematic showing the gRNA design, filtering, and annotation strategy (see Star Methods). Each human chromosome harbours the (peri)centromeric region (Pan-centromere), active alpha-satellite higher order repeat region (Active αSat HOR), and CENPA-bound region (CENPA-bound) in the hierarchical structure shown. The gRNA design uses the CENPA-bound sequences from T2T-CHM13v2.0 reference, resolved and annotated by the T2T consortium. The gRNA filtering considers gRNA GC-content (35% – 80%), the number of binding sites (≥20), and CENPA-bound chromosome specificity (≥ 0.9). Three additional specificity scores are calculated and annotated for the 1830 filtered chromosome-specific gRNAs. 15 bi-chromosome-specific gRNAs targeting Chr13/21 pair are designed. The gRNA information is summarised in Table S1. (B) Schematics showing the screening setup for identifying functional gRNAs that induce chromosome-specific micronucleated cell formation. Cells are electroporated with dCas9-gRNA RNP complex and cultured for 48 h. DAPI staining is performed to quantify the percentage of micronucleated cells (efficiency). Fluorescence *in situ* hybridization (FISH) is performed to validate the identity of the chromosome within the micronucleus. The bottom table indicates the FISH probes used for each chromosome – ‘Cen’ represents the centromeric probe and ‘pTel’ or ‘qTel’ represents the telomeric probe specific to chromosome p-arm or q-arm, respectively. The gRNA screening results are in Table S1.

**Figure S2.**
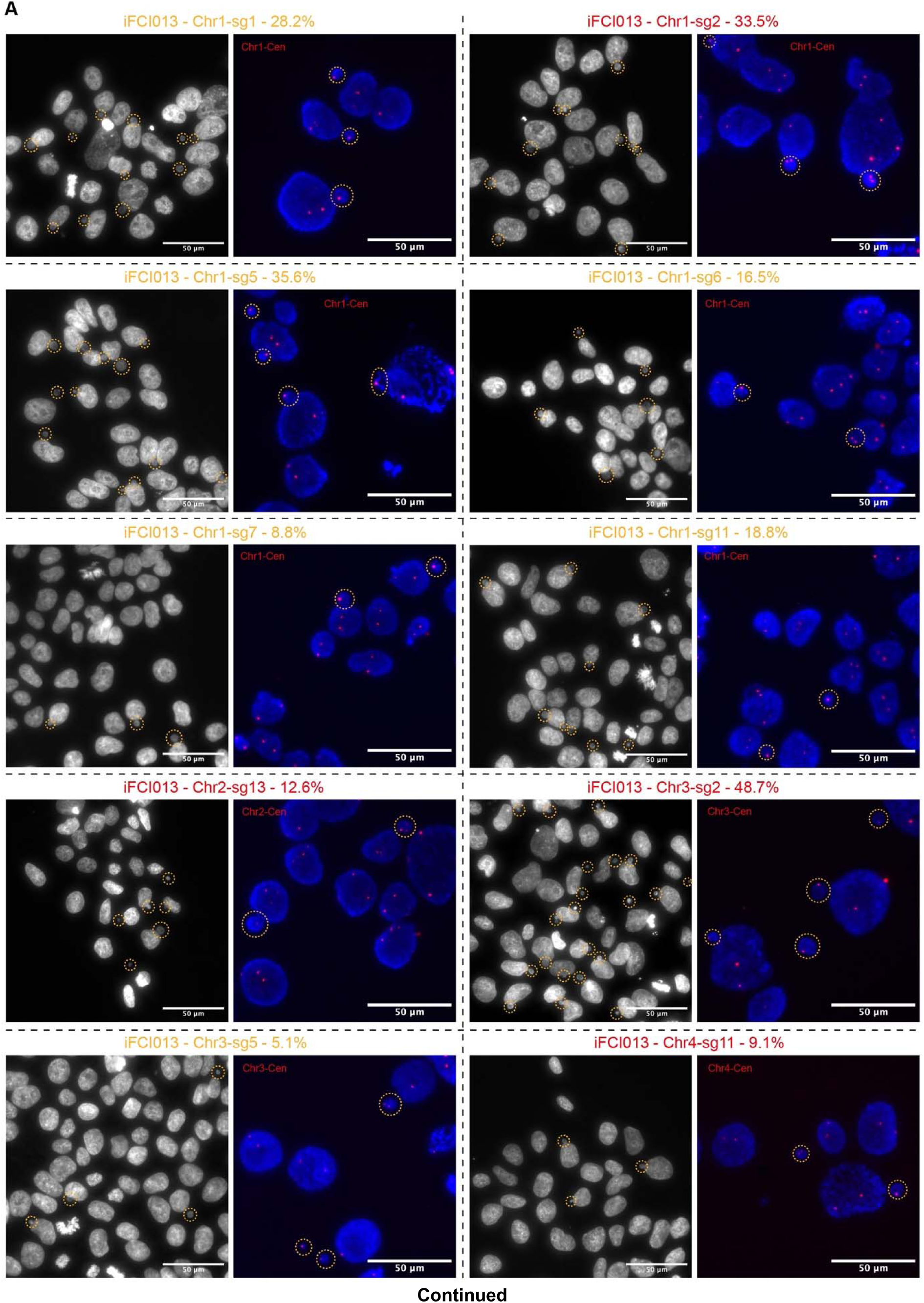
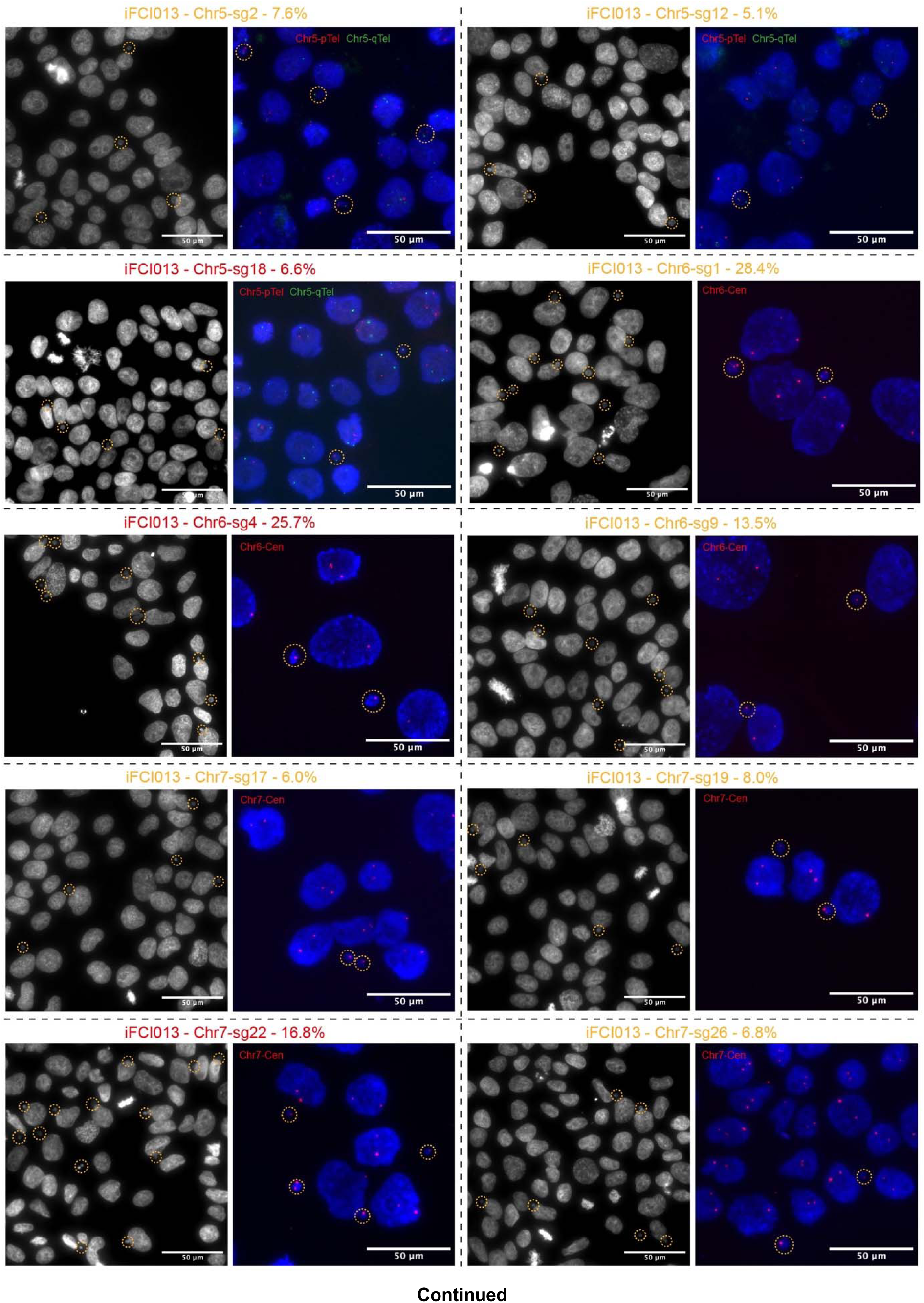
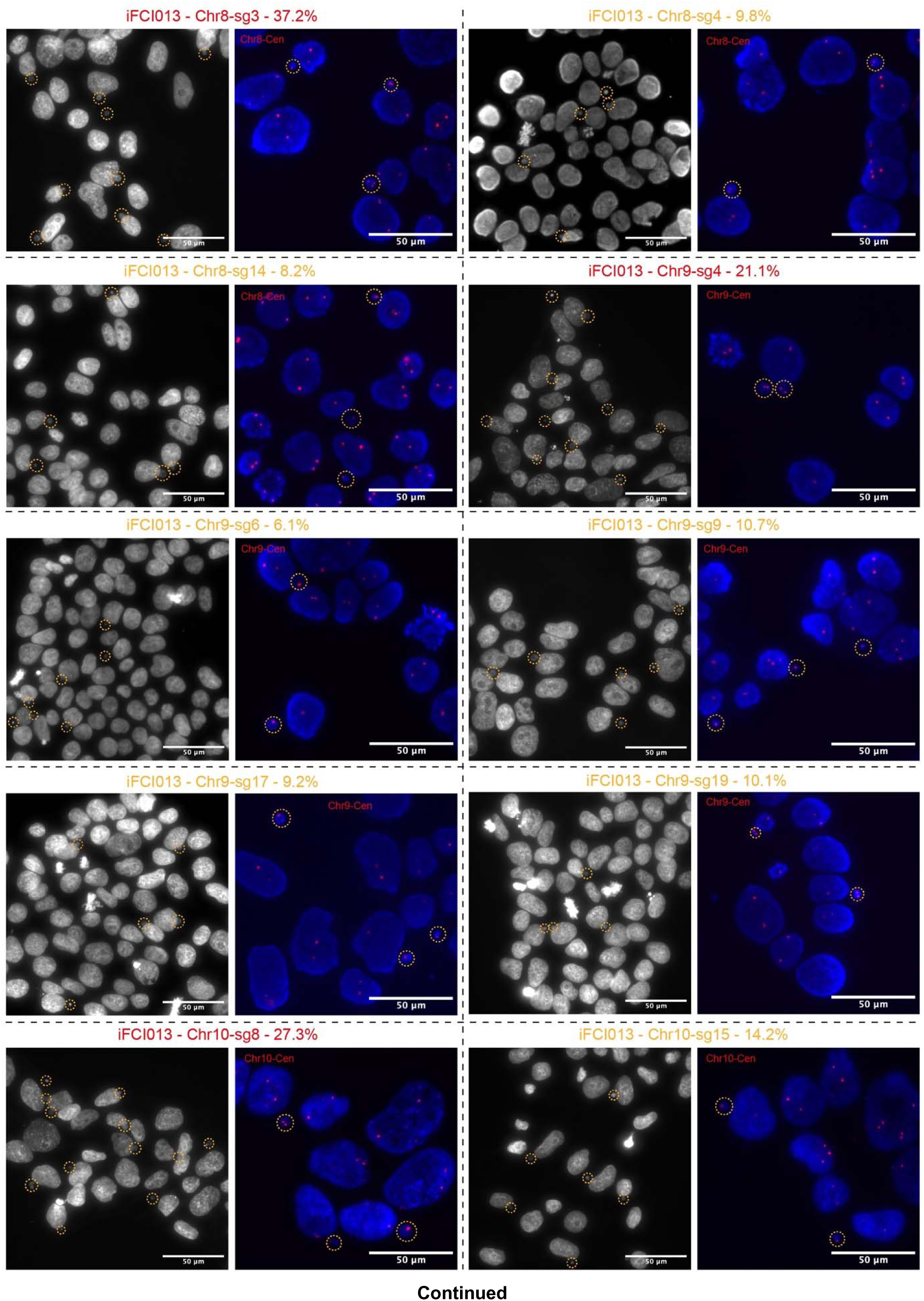
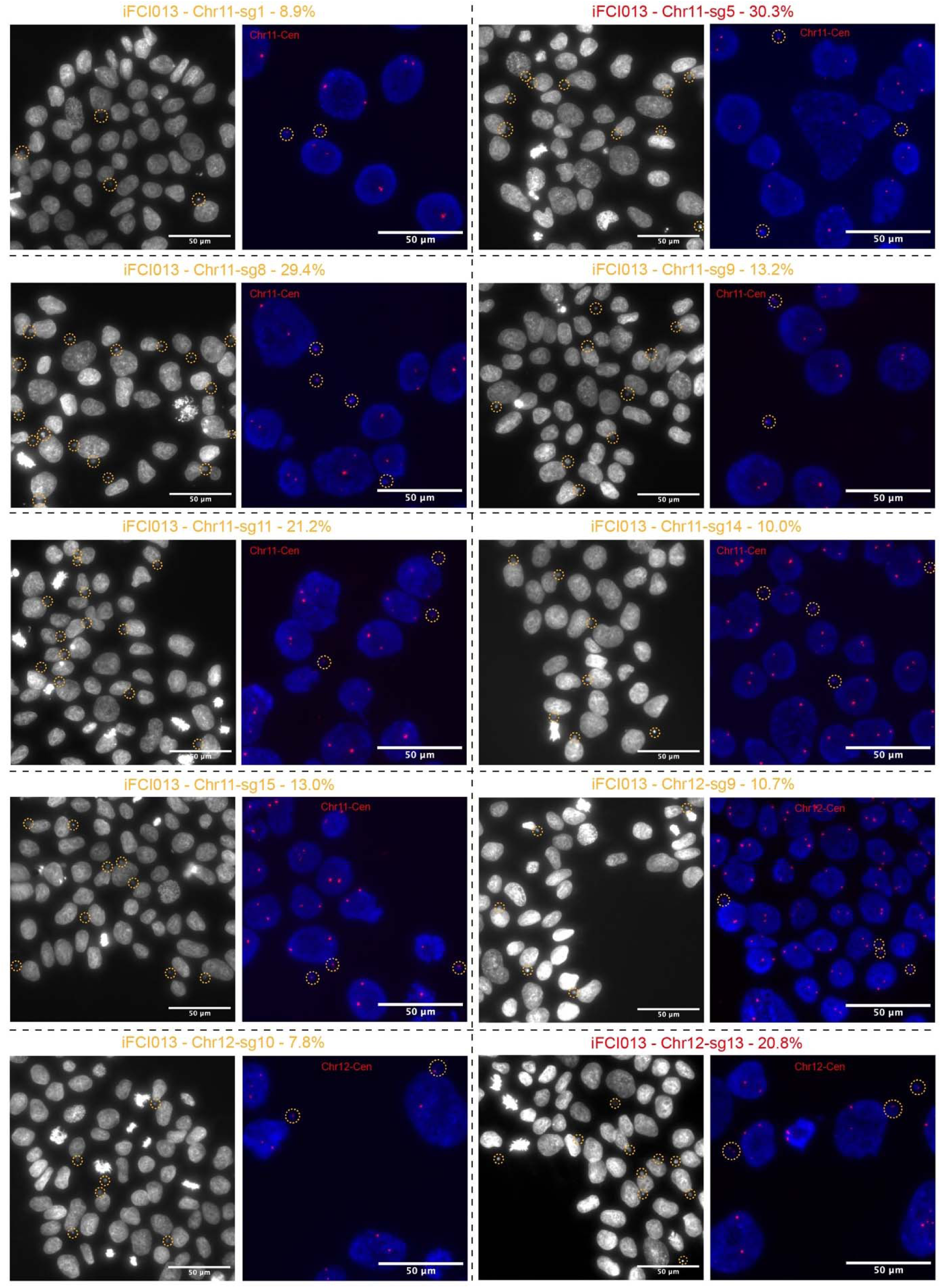
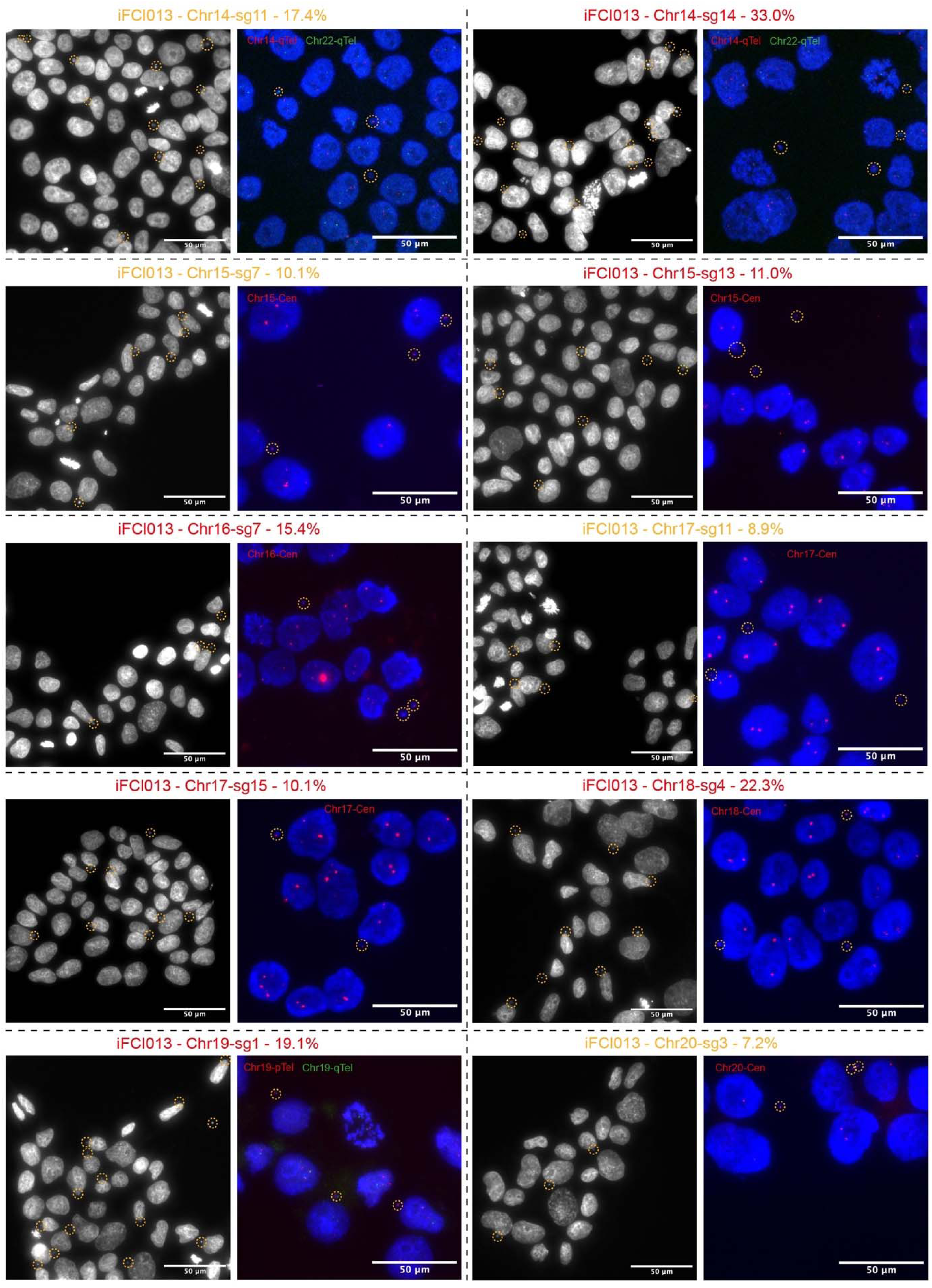
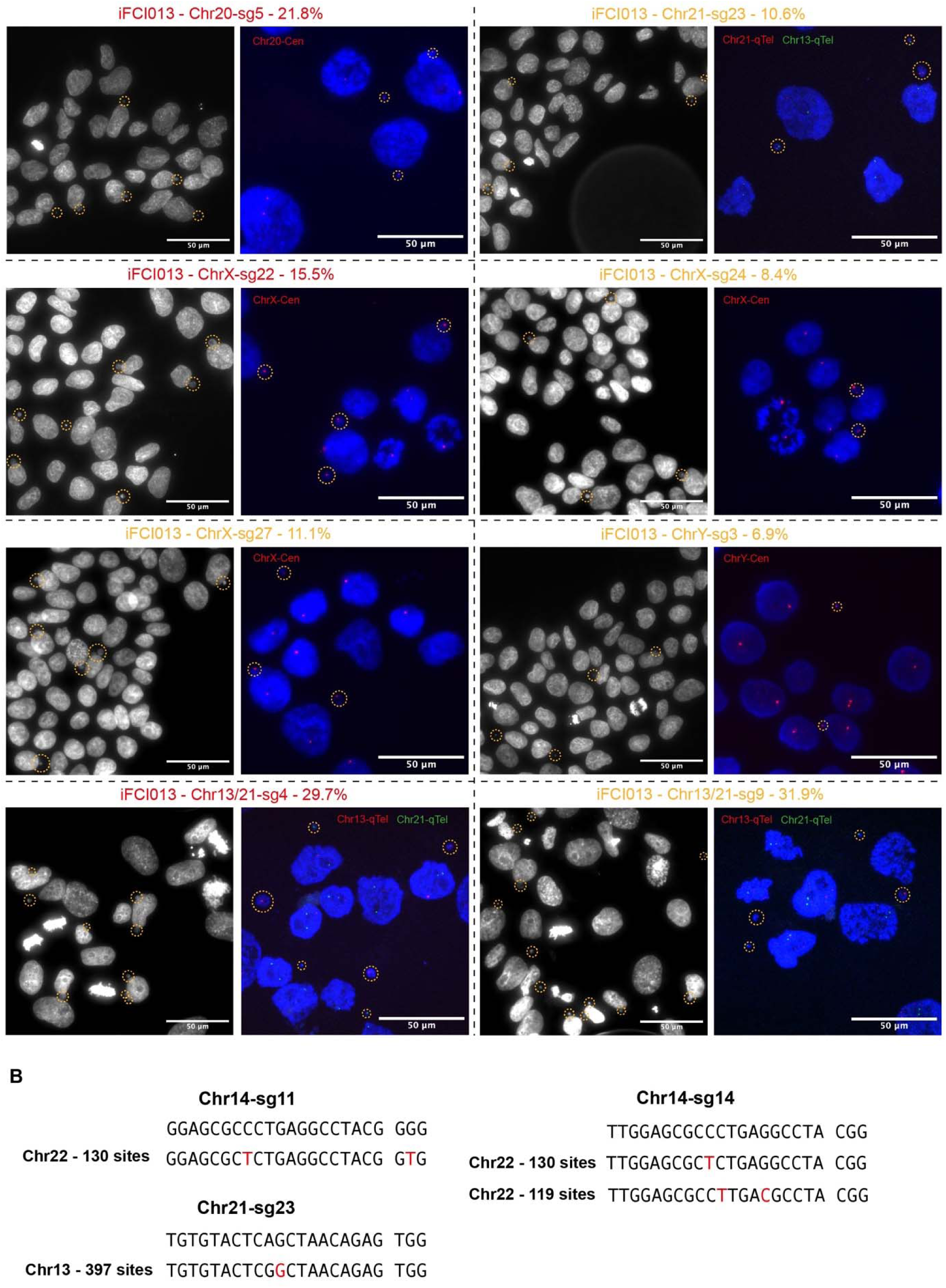
Functional CRISPR-Taiji gRNAs induce chromosome-specific micronucleus formation in iFCI013 iPSCs, related to Figure 1. (A) Representative images showing the micronucleus induction results of the 58 identified functional gRNAs in iFCI013 iPSCs. For each gRNA, the percentage of micronucleated cells at 48 h from the initial gRNA screen is annotated next to the gRNA ID and the selected gRNAs for further validations are shown in red; the left image shows the micronucleated cell formation with DAPI staining (micronuclei circled in dashed line), and the right image shows FISH validation of chromosome identity within micronucleus (circled in dashed line) with the indicated FISH probe(s). Scale bars, 50 μm. (B) DNA sequence alignments showing the predicted off-target gRNA binding sequences with more than twenty binding sites and no more than two mismatches (shown in red) at the CENPA-bound region of the observed off-target chromosomes for the indicated functional gRNAs. The off-target chromosome and the number of predicted off-target sites are shown.

**Figure S3.**
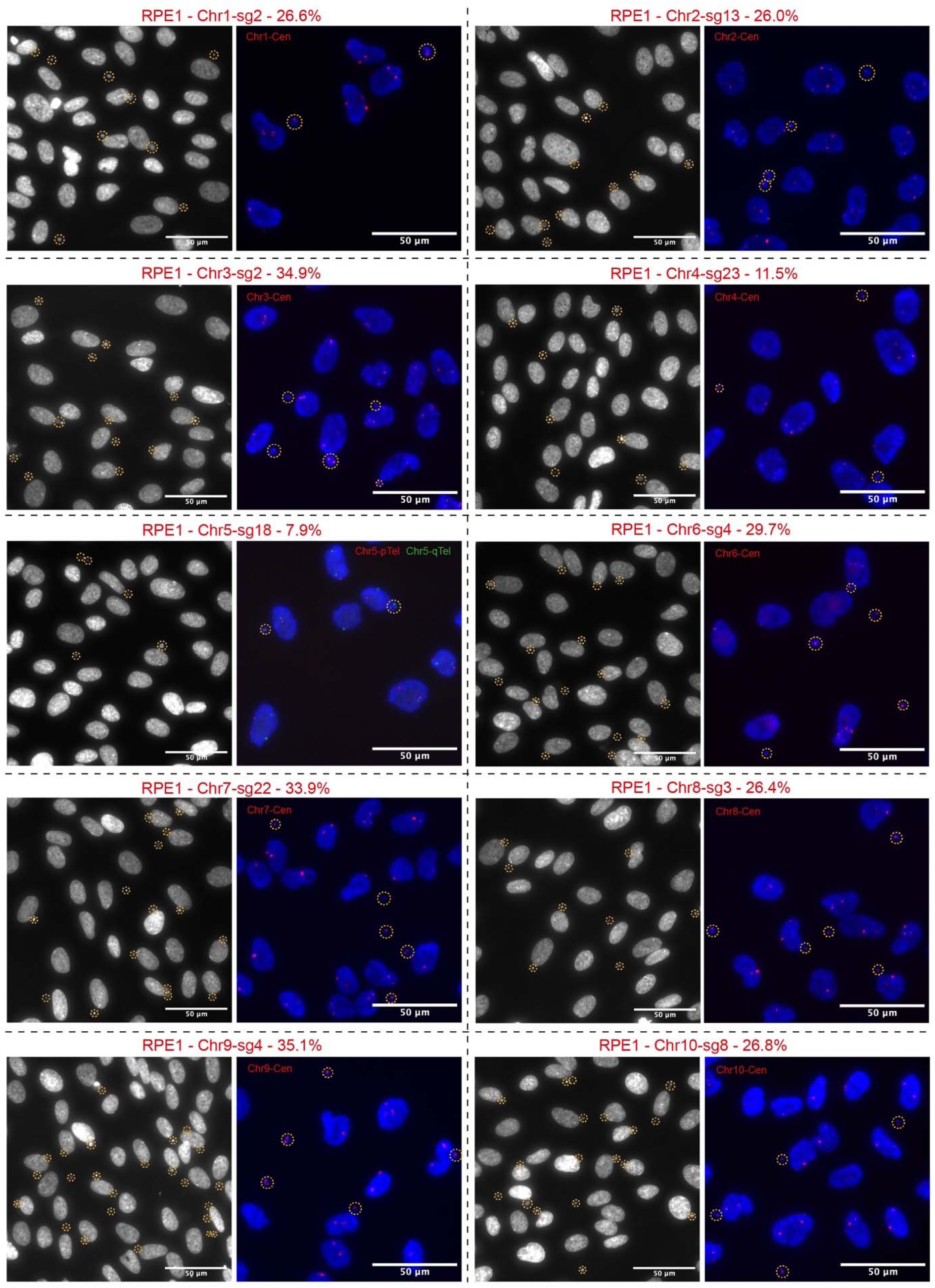
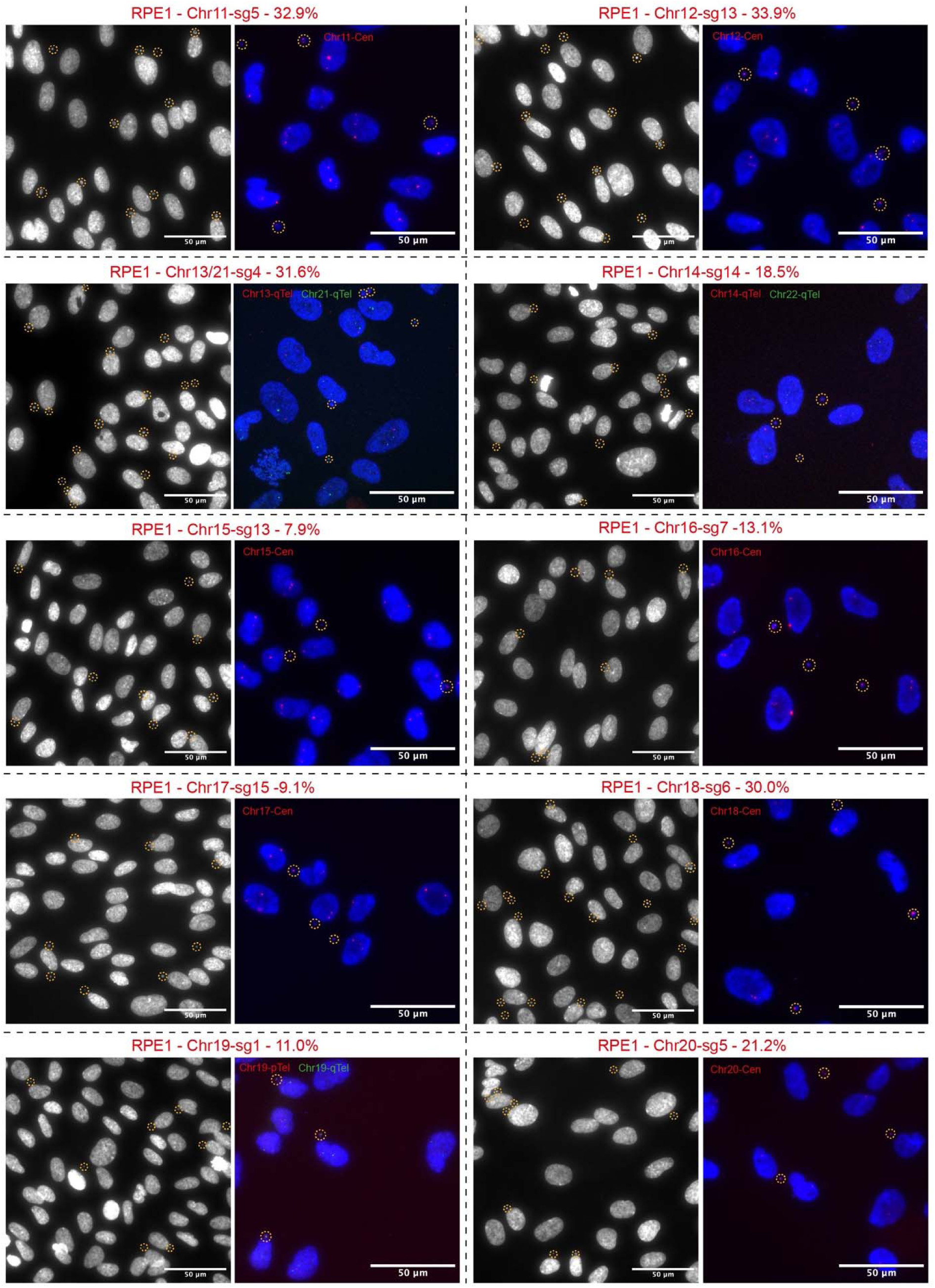
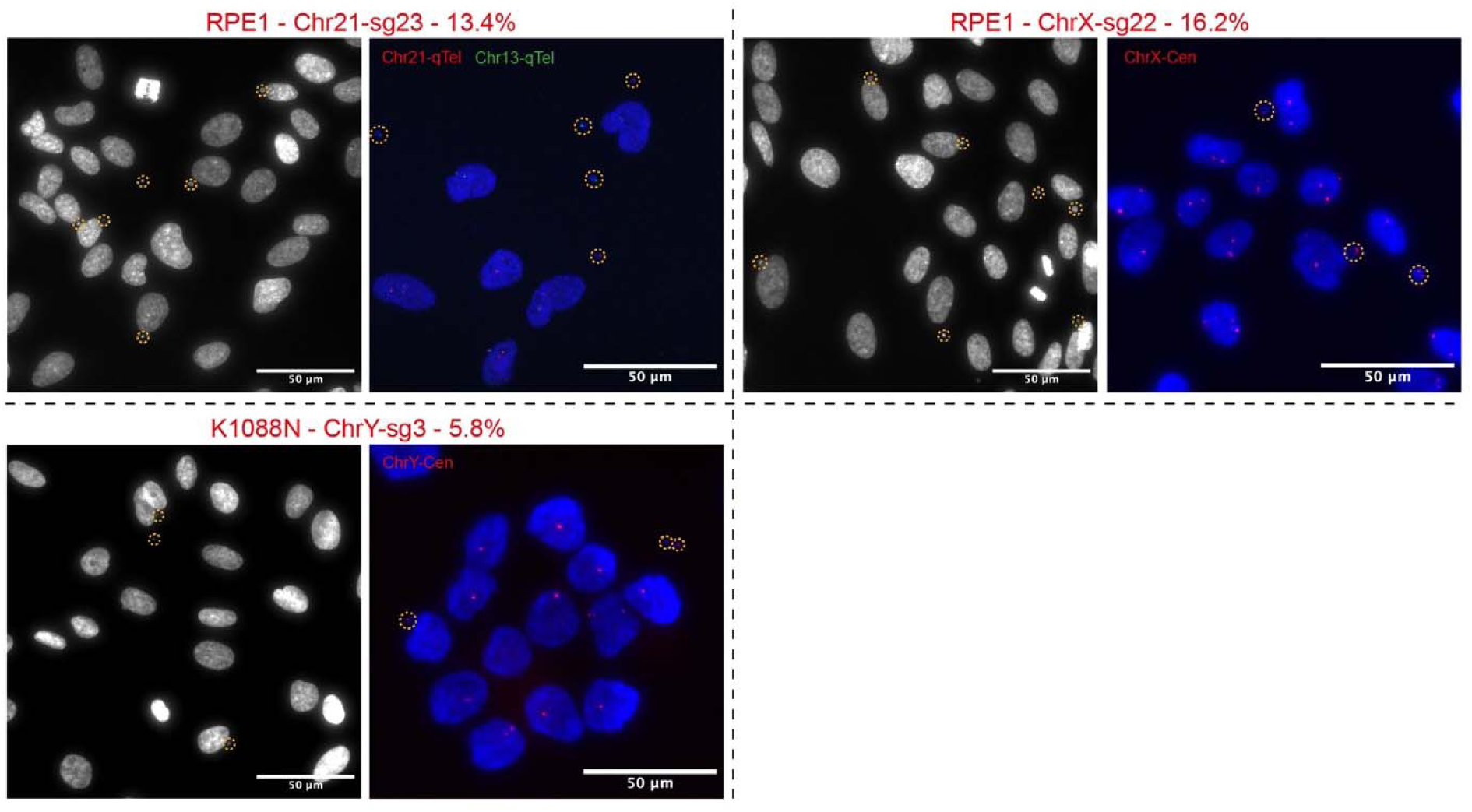
Validation of the selected functional gRNAs in RPE1 or K1088N cells, related to Figure 1. Representative images showing the characterization results of the 23 selected functional gRNAs in RPE1 or K1088N cells. For each gRNA, the cell line used and the average percentage of micronucleated cells at 48 h of three independent repeats is annotated next to the gRNA ID; the left image shows the micronucleated cell formation with DAPI staining (micronuclei circled in dashed line), and the right image shows FISH validation of chromosome identity within micronucleus (circled in dashed line) with the indicated FISH probe(s). Scale bars, 50 μm.

**Figure S4.**
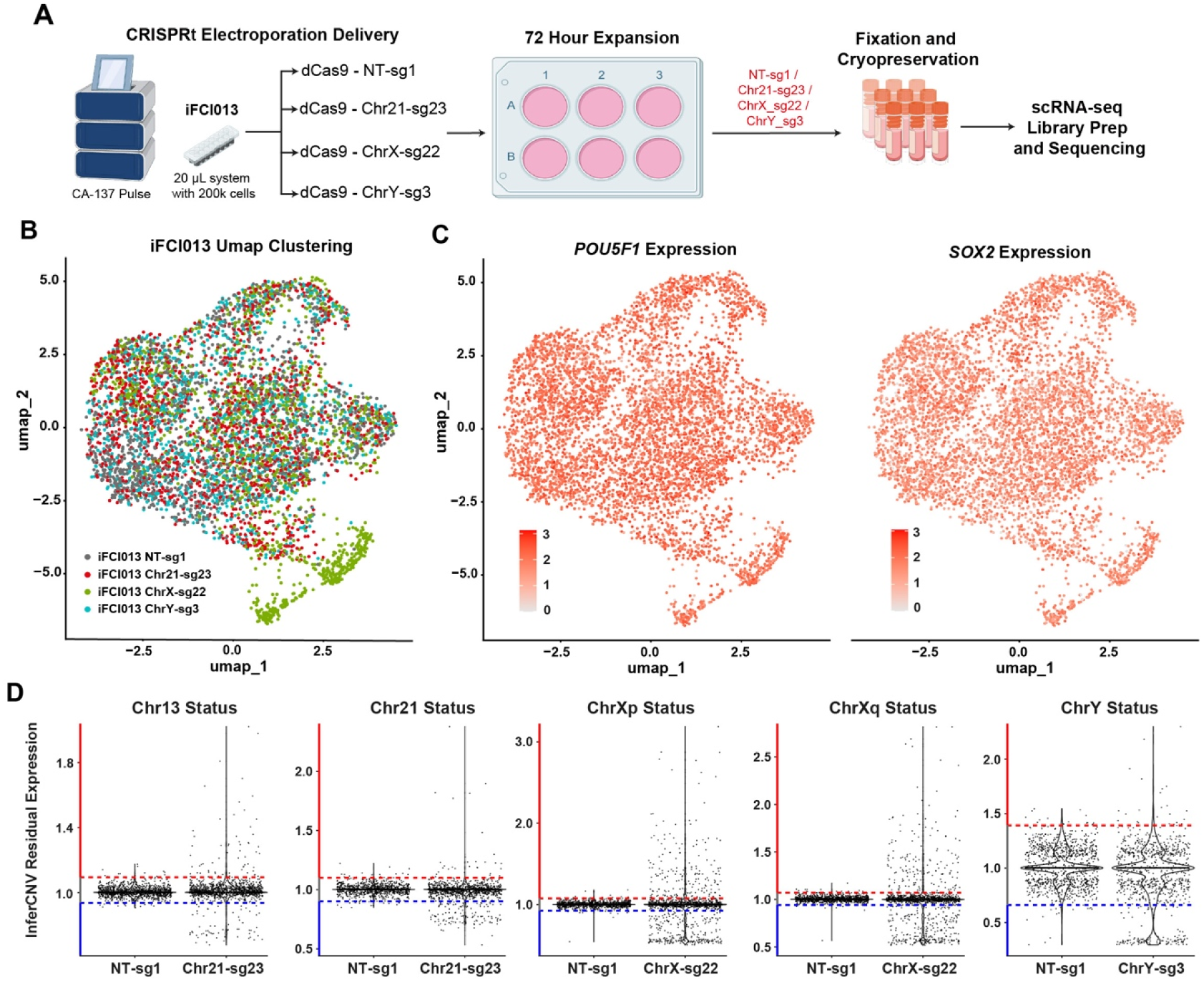
Chr13, 21, X or, Y CRISPRt aneuploidy engineering in iFCI013 iPSCs, related to Figure 1. (A) Schematics showing the experiment setup of Chr13/21, X, or Y aneuploidy engineering using CRISPRt with the indicated gRNAs in iFCI013 cells. The four samples fixed for scRNA-seq are marked in red. (B) UMAP clustering plot of the scRNA-seq analysis in iFCI013 cells. (C) Feature plots showing the expression of pluripotency genes *POU5F1* (OCT4) and *SOX2* in individual iFCI013 cells. Color gradient represents the normalized expression levels. (D) Violin plots showing InferCNV residual expression across the indicated chromosomes or chromosome arms after the indicated CRISPRt. The dots represent individual cells. The blue and red dashed lines represent the thresholds (see Star Methods) for copy number loss and gain, respectively. The cell proportions with the indicated copy number status are shown in the tables.

**Figure S5.**
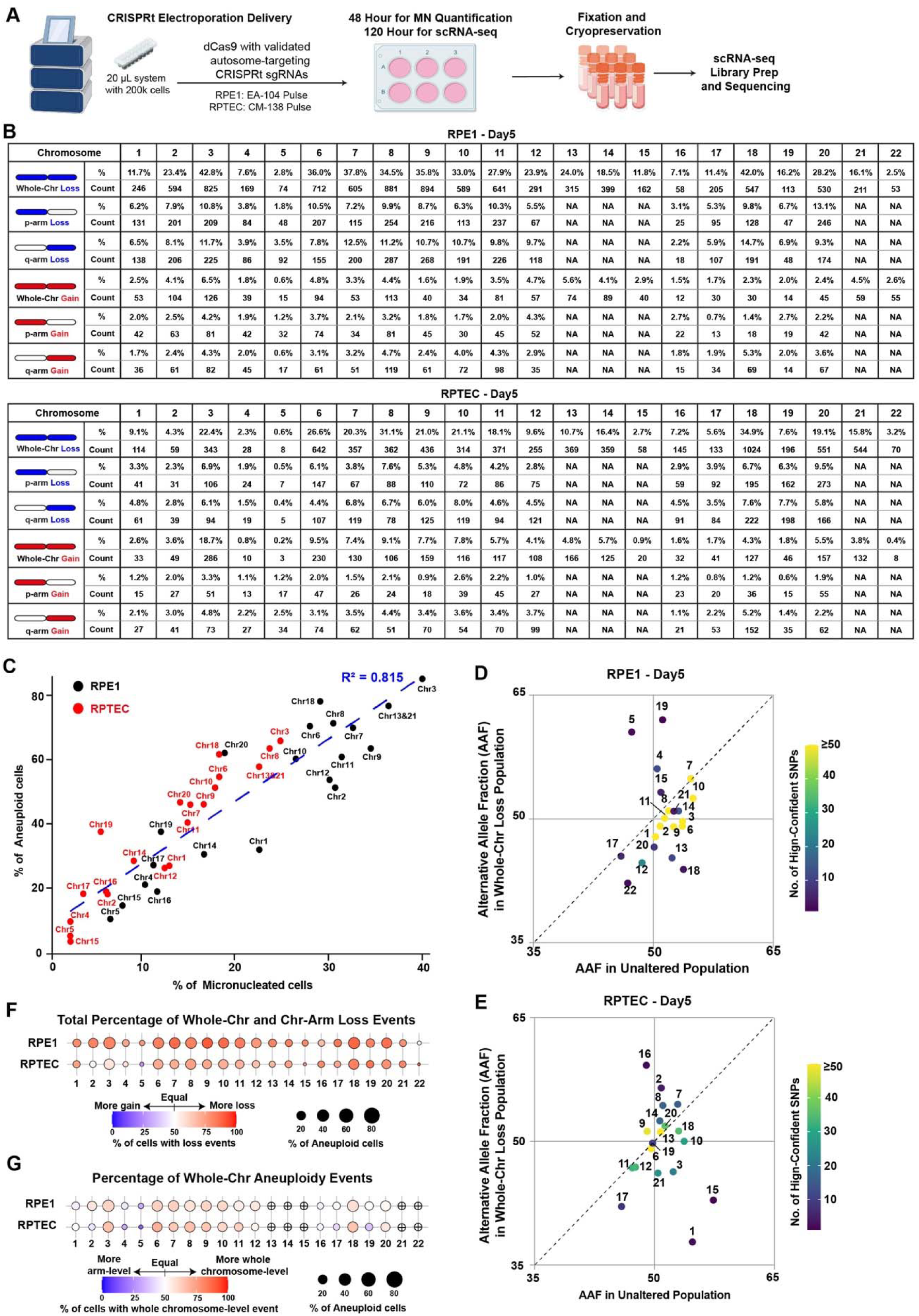
Pan-autosome CRISPRt aneuploidy engineering in RPE1 and RPTEC cells using CRISPRt, related to Figure 2. (A) Schematics showing the experiment setup of pan-autosome (Chr1-22) aneuploidy engineering using CRISPRt with the validated gRNA panels in RPTEC cells. (B) Summary table showing the frequencies of indicated whole-chr or chr-arm aneuploidy events in RPE1 and RPTEC CRISPRt experiments (Figure 2). (C) Scatter plot showing the correlation between the percentages of micronucleated (x-axis) and aneuploid (y-axis) cells in RPE1 (black) and RPTEC (red) for indicated CRISPRt experiments (Figure 2). The linear regression line and Pearson’s correlation coefficient square value (R^2^) are shown. (D and E) Scatter plot showing the average alternative allele frequencies of single-nucleotide polymorphisms (SNPs) detected in scRNA-seq data across the indicated chromosomes in RPE1 (D) and RPTEC (E) cells following CRISPRt aneuploidy engineering. Each dot represents one chromosome, with the x– and y-axes representing cell populations with the classified unaltered (copy number neutral) and whole-chr loss statuses, respectively. The dot color represents the number of high-confident SNPs included in the quantification. (F and G) Dot plots showing the percentages of cells with chromosomal loss aneuploidy events (F) and whole-chr aneuploidy events (G) among all aneuploidy cells for each indicated chromosome in RPE1 and RPTEC cells following CRISPRt aneuploidy engineering.

**Figure S6.**
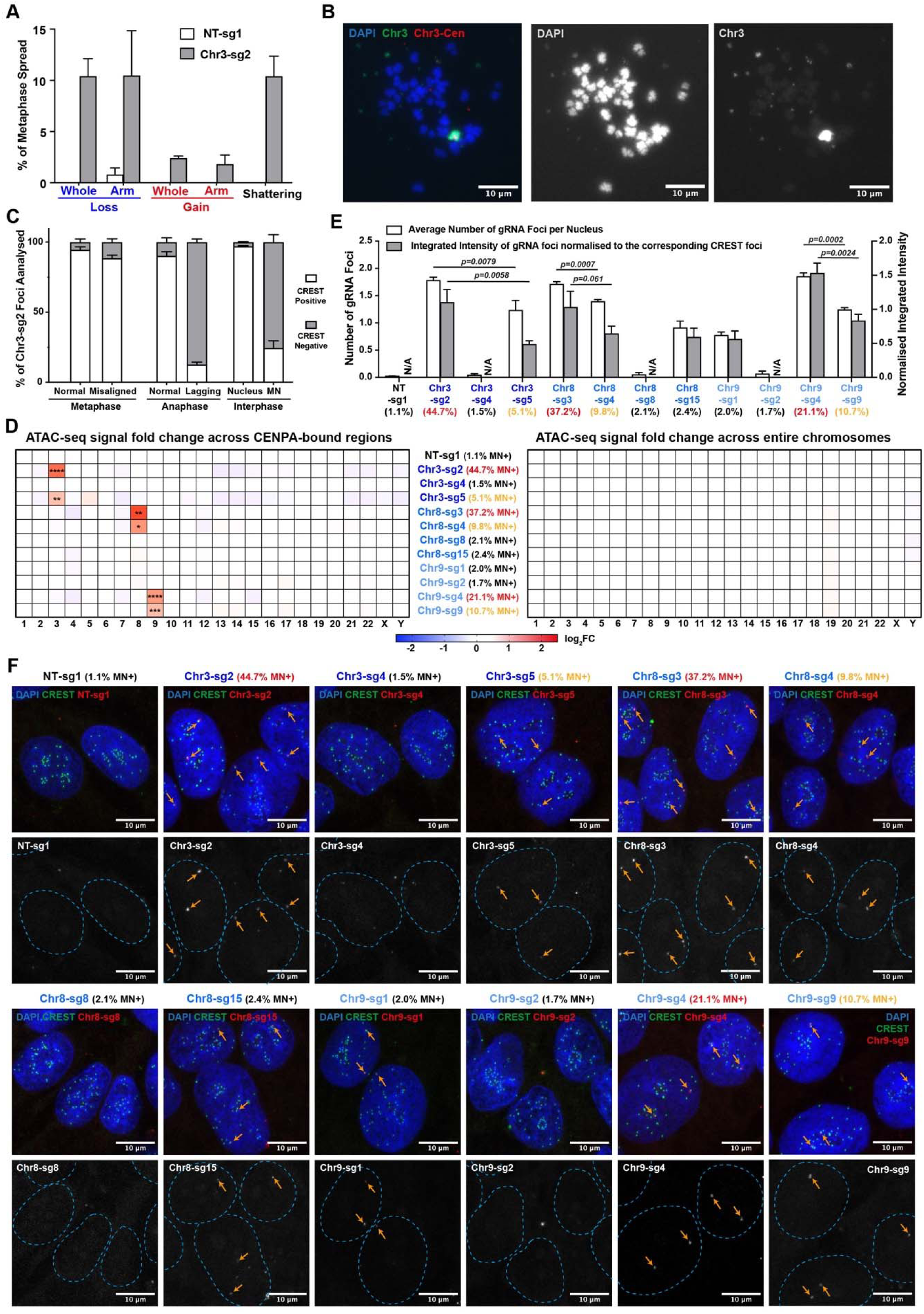
CRISPRt generates a diverse spectrum of chromosomal alterations via dCas9-induced centromeric chromatin relaxation, related to Figure 3. (A) Bar plot showing percentage of indicated Chr3 aberrations quantified by metaphase spread and chromosome painting in RPTEC-shTP53 cells 7 days following CRISPRt aneuploidy engineering. Error bars, mean ± SD (n=2, >400 metaphase spreads). (B) Representative images showing shattered Chr3 fragments 7 days in RPTEC-shP53 cells following CRISPRt aneuploidy engineering. Scale bars,10 μm. (C) Bar plot showing percentage of quantified Chr3 centromeres (Chr3-sg2 foci) with or without inner kinetochore components (CREST positive or negative) on Chr3 with the indicated normal and abnormal status in metaphase, anaphase and interphase 24 h after Chr3 CRISPRt in iFCI013 cells (representative images in Figure 2D-F). (n=3, >100 Chr3-sg2 foci quantified for each status). (D) Heatmaps showing the average log_2_ fold changes (log_2_FC) of ATAC-seq signal in iFCI013 cells 8 h after CRISPRt with the indicated gRNAs compared to NT-sg1, across the CENPA-bound regions only (left) or across the entire chromosomes (right). (n=2, representative tracks in Figure 3G, one-way ANOVA followed by Dunnett’s post hoc multiple comparison test compared to NT-sg1 across individual chromosomes, significant changes are shown as ****p<0.0001, ***p<0.001, **p<0.01, and *p<0.05). (E) Bar plot showing average number of centromeric dCas9-gRNA foci per nucleus and average normalized intensity dCas9-gRNA foci (normalized to the corresponding overlapped CREST foci intensity, see Star Methods) 8 h after CRISPRt with the indicated gRNAs in iFCI013 cells. The micronucleated cell induction efficiencies of gRNAs are shown. Error bars, mean ± SD (n=3, >300 nuclei quantified, two-tailed t-test with p values shown). (F) Representative images showing the centromeric recruitment of dCas9 by different CRISPRt gRNAs with indicated micronucleated cell induction efficiencies 8 h after CRISPRt in iFCI013 cells. CREST antibody marks the centromere, and Atto-550 labelled gRNA marks dCas9. Arrows highlight the centromeric dCas9-gRNA foci. Scale bars,10 μm.

**Figure S7.**
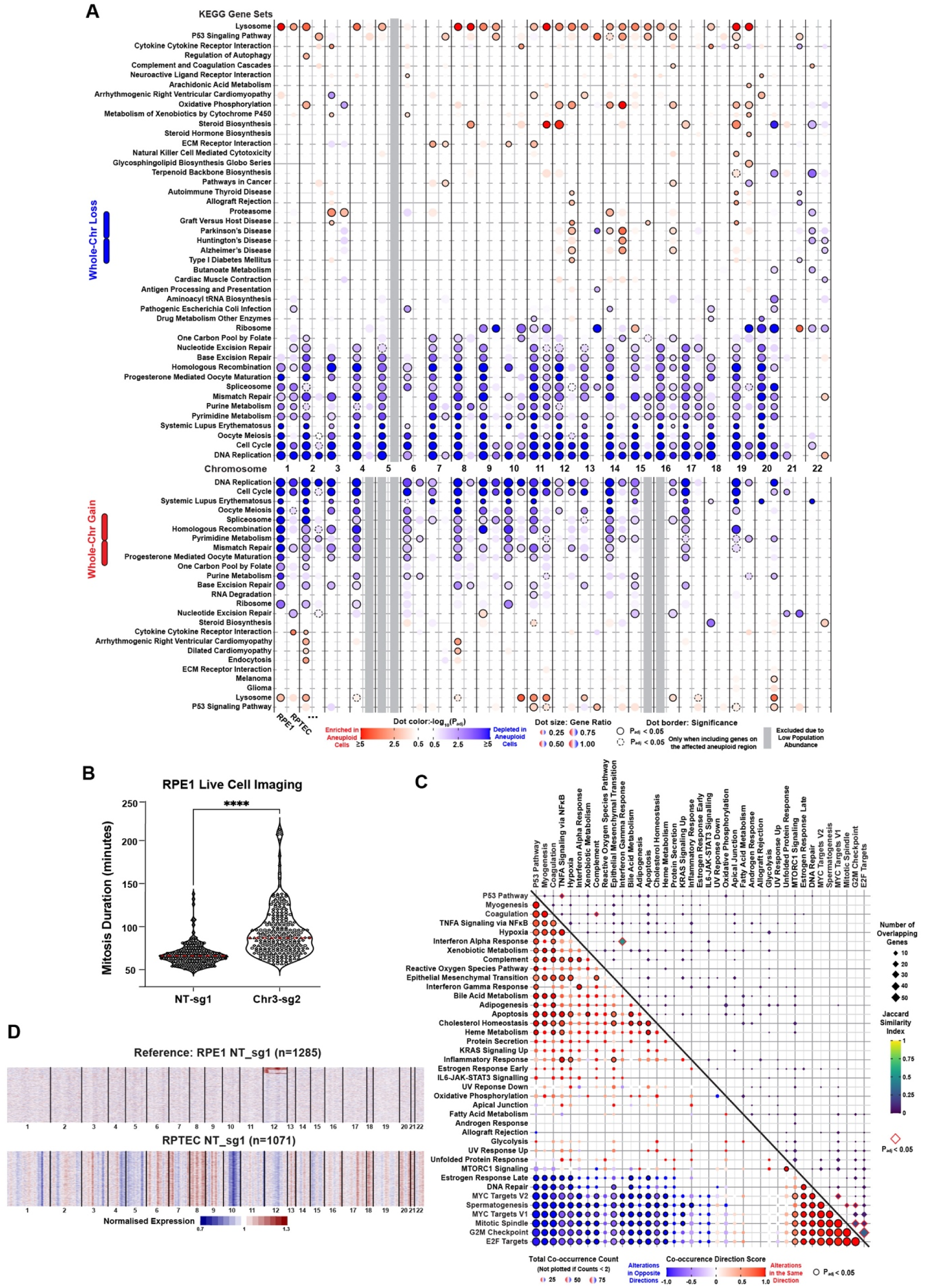
Transcriptomic analysis reveals universal and chromosome-specific alterations across whole-chr aneuploidies, related to Figure 4. (A) Dot plot showing significantly enriched or depleted KEGG gene sets by GSEA in RPE1 and RPTEC cells with the indicated whole-chr aneuploidy, compared to the corresponding copy-neutral population. The main analyses exclude genes located within the corresponding aneuploid region to highlight the secondary genome-wide effects. The columns represent cell type and chromosomes. The rows represent significantly altered KEGG gene sets, ordered by average enrichment scores. The color gradient represents –log_10_(P_adj_) values with red and blue corresponding to positive and negative normalized enrichment scores, respectively. Dots with a solid border highlight significant changes (P_adj_ <0.05). Dots with a dashed border highlight additional significant changes when all the genes are included in the analyses for the global changes. (B) Violin plot showing the mitotic duration by time-lapse live-cell imaging after CRISPRt with the indicated gRNAs in RPE1 cells. (n=3, >200 mitotic events quantified). (C) Dot plot showing co-occurrence of significant GSEA hallmark gene set alterations (bottom left triangle) and overlaps between GSEA leading-edge gene subsets (upper right triangle) in RPE1 and RPTEC cells with whole-chr aneuploidies (related to Figure 4). For co-occurrence, each dot correlates to a pair of gene sets. The dot size represents the total co-occurrence count (the number of times when a pair of gene sets significantly altered in the same aneuploid population). The dot color gradient represents the co-occurrence direction score, with red indicating co-occurrence biased towards the same direction (both enriched or both depleted) and blue indicating co-occurrence biased towards opposite directions (one enriched, one depleted). Dots with black borders indicate statistically significant co-occurrence directionality (Padj < 0.05, two-sided binomial test with Benjamini-Hochberg procedure). For the overlaps between leading-edge genes, each diamond correlates to a pair of gene sets. The diamond size represents the number of overlapping genes in the leading-edge gene subsets. The dot color gradient represents the Jaccard similarity index. Diamonds with red borders indicate statistically significant overlaps (Padj <0.05, Fisher’s exact test with Benjamini-Hochberg procedure). (D) Heatmaps showing the global transcriptomic difference between the RPE1 (as reference) and RPTEC cells, quantified from scRNA-seq data. Each row represents an individual cell; columns correspond to chromosomes, scaled by the number of expressed genes. The consistently lower expression of Chr10q genes in RPTEC cells reflects the clonal Chr10q gain present in RPE1 cells, which is treated as copy number neutral in the RPE1 NT-sg1 population as the reference.

**Figure S8.**
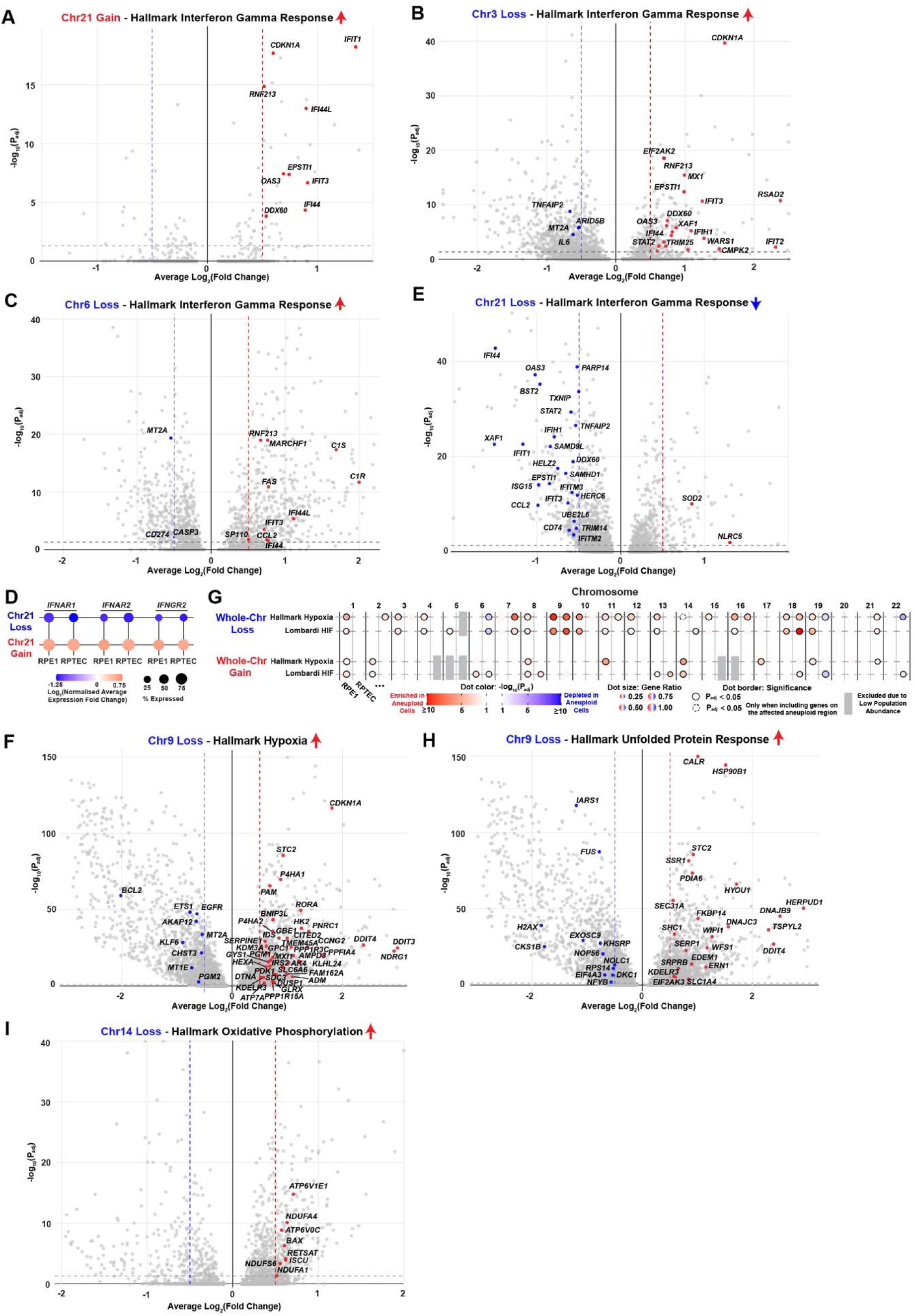
Specific whole-chr aneuploidies show lineage-conserved transcriptomic alterations, related to Figure 4. (A, B, C and E) Volcano plots showing differentially expressed genes (DEGs) in engineered Chr21 gain (A), Chr3 loss (B), Chr6 loss (C), and Chr21 loss (E) populations, compared to the corresponding copy-neutral cell population. Significant DEGs (Padj<0.05, Log_2_(Fold Change)>0.5 or <-0.5) within hallmark interferon gamma response gene sets are labelled with HGNC symbols in the corresponding figures. (D) Dot plot showing the average normalized expression changes of interferon receptor genes located on Chr21 in RPE1 and RPTEC cell populations with Chr21 loss or gain engineered by CRISPRt, compared to the corresponding copy-neutral cell populations. (F, H and I) Volcano plots showing differentially expressed genes (DEGs) in engineered Chr9 loss (F and H) and Chr14 loss (I) populations, compared to the corresponding copy-neutral cell population. Significant DEGs (Padj<0.05, Log_2_(Fold Change)>0.5 or <-0.5) within hallmark hypoxia (F), unfolded protein response (H), and oxidative phosphorylation (I) gene sets are labelled with HGNC symbols in the corresponding figures. (G) Dot plot comparing the enrichment or depletion of hallmark hypoxia gene set and a stringent hypoxia inducible factor (HIF) target gene set^37^ (Lombardi HIF) in RPE1 and RPTEC cells with the indicated whole-chr aneuploidy, compared to the corresponding copy-neutral population. The analyses exclude genes located within the corresponding aneuploid region to highlight secondary genome-wide effects. The columns represent cell type and chromosomes. The color gradient represents –log_10_(P_adj_) values with red and blue corresponding to positive and negative normalized enrichment scores, respectively.

**Figure S9.**
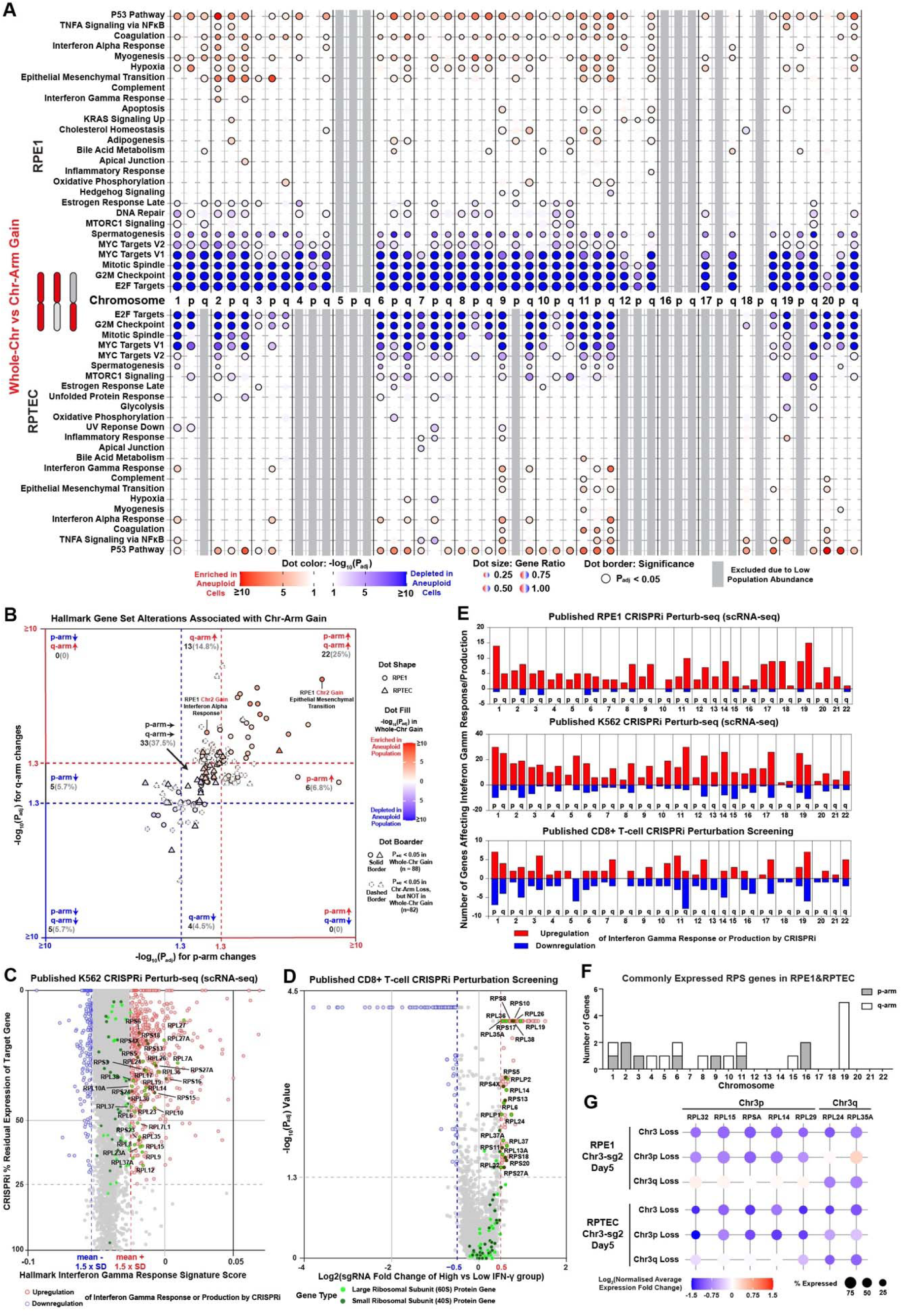
Chr-arm and gene-level changes likely underlie specific alterations associated with whole-chr aneuploidies, related to Figure 5. (A) Dot plot showing significantly enriched or depleted hallmark gene sets by GSEA in RPE1 and RPTEC cells with indicated whole-chr and chr-arm gains, compared to the corresponding copy-neutral population. The analyses exclude genes located within the corresponding aneuploid region to highlight secondary genome-wide effects. The columns represent the indicated whole-chr or chr-arm gain. The color gradient represents –log_10_(P_adj_) values with red and blue corresponding to positive and negative normalized enrichment scores, respectively. Dots with a solid border highlight significant changes (P_adj_ <0.05). (B) Scatter plot showing the hallmark gene set alterations associated with the corresponding chr-arm gain in RPE1 (circles) and RPTEC (triangles) cells, excluding the universal alterations. Each dot represents an altered gene set associated with whole-chr or chr-arm gain of a specific chromosome in RPE1 or RPTEC cells, with highlighted gene sets labelled by name, cell type and chromosome. The x– and y-axes show the –log_10_(P_adj_) values for the corresponding p-arm and q-arm gains, respectively, with color indicating the direction of enrichment. Dot fill color shows the –log_10_(P_adj_) values and the direction of enrichment observed in the corresponding whole-chr gain. Transparent dots with dashed borders denote gene sets that are significantly altered only in the chr-arm gain. The number and percentage of significant gene set alterations in the whole-chr gain with the indicated changes in the corresponding p-arm and q-arm gains are shown. (C) Scatter plot showing the correlation between CRISPRi-induced gene perturbations and hallmark interferon gamma response signature scores with the published genome-scale Perturb-seq dataset in K562 cells^29^. The x– and y-axes show the signature score and the residual expression of the target gene after CRISPRi knockdown. Ribosomal large subunit protein (RPL) and small subunit protein (RPS) genes with a significant effect on interferon gamma response are labelled. (D) Volcano plot showing the effects of CRISPRi-induced gene perturbation on interferon gamma production with the published genome-scale Perturb-seq dataset in primary CD8⁺ T cells^44^. The x-axis indicates log₂ fold change of gRNA enrichment in interferon gamma high versus low populations; the y-axis shows the corresponding –log_10_(P_adj_) values. RPL and RPS genes with a significant effect on interferon gamma production are labelled. (E) Bar plots showing the number of genes whose CRISPRi knockdown significantly alters interferon gamma response signature score with the published essential-wide Perturb-seq dataset in RPE1^29^ (top) and the genome-wide Perturb-seq dataset in K562 cells^29^ (middle), or interferon gamma production with the published genome-wide Perturb-seq dataset in primary CD8⁺ T cells^44^ (bottom). Genes with CRISPRi knockdown resulting in upregulation (red) or downregulation (blue) of interferon gamma signature/production are quantified based on their chromosomal locations. (F) Histogram showing the distribution of consistently expressed RPS genes in RPE1 and RPTEC cells across chromosome arms. (G) Dot plot showing the average normalized expression changes of RPL and RPS genes located on Chr3 in RPE1 and RPTEC cell populations with whole-Chr3 or Chr3p-arm or Chr3q-arm loss engineered by CRISPRt, compared to the corresponding copy-neutral cell populations.

**Figure S10.**
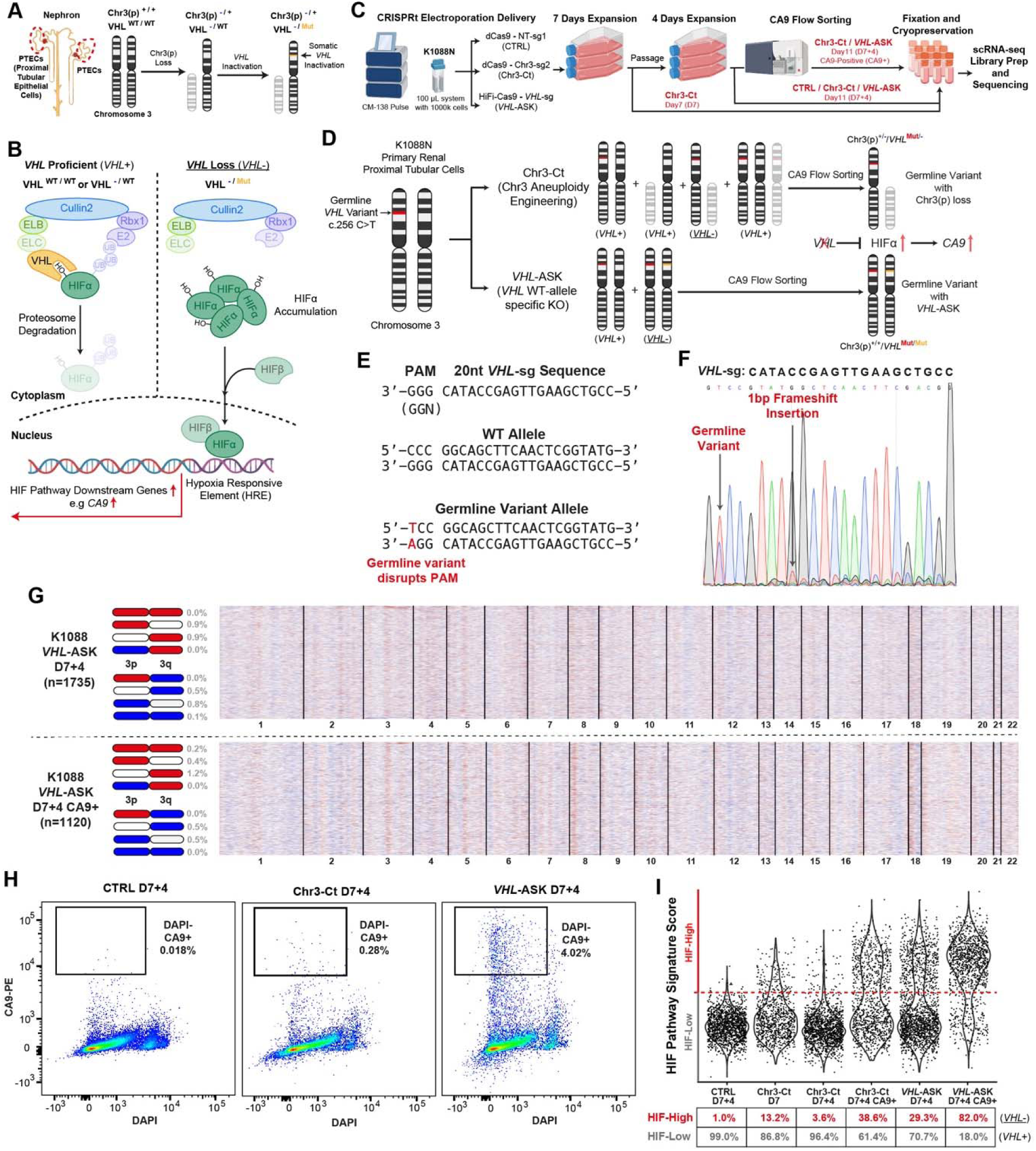
Engineering and enrichment of Chr3(p) loss population in K1088N PTECs, related to Figure 6. (A) Schematics showing the tumor initiation from normal renal proximal tubule epithelial cells (PTECs) for sporadic ccRCC through Chr3(p) loss and *VHL* somatic inactivation. Chr3(p) allele status is annotated as ‘+’ (unchanged) or ‘-’ (loss). *VHL* gene status is annotated as ‘WT’ (wild-type), ‘Mut’ (mutated), or ‘-’ (deleted). (B) Schematics showing *VHL* protein function in repressing the hypoxia-inducible factor (HIF) signaling pathway when *VHL* is proficient (*VHL*+, at least one WT *VHL* copy) under normal oxygen conditions, and the aberrant constitutive HIF pathway activation when *VHL* is lost (*<u>VHL</u>*<u>-</u>, both *VHL* copies inactivated), driving expression of HIF pathway downstream genes. (C) Schematics showing the experiment setup of Chr3 aneuploidy engineering using CRISPRt with Chr3-sg2 (Chr3-Ct) or NT-sg1 (CTRL), and *VHL* allele-specific KO (*VHL*-ASK) in K1088N PTECs. Cells were cultured for 7 days (D7) or 11 days (D7+4) after treatments. Cells with Chr3-Ct or *VHL*-ASK on D7+4 were selected for CA9-positive (CA9+) cells with flow sorting. The six samples fixed for scRNA-seq are marked in red. (D) Schematics showing the expected genotypes generated after Chr3-Ct or *VHL*-ASK in K1088N PTECs with the corresponding *VHL* status annotated (*VHL*+ for *VHL* proficient, and *<u>VHL-</u>* for *VHL* loss) and the target genotypes with *VHL* loss enriched by CA9-positive (CA9+) flow sorting. (E) Schematics showing the design of gRNA (*VHL*-sg) for *VHL* allele-specific KO (*VHL*-ASK) in K1088N PTECs. The *VHL* germline variant in K1088N cells disrupts the protospacer adjacent motif (PAM) of *VHL*-sg gRNA. (F) Sanger sequencing chromatogram showing the 1bp frameshift insertion generated by *VHL*-ASK in K1088N PTECs. (G) Heatmaps showing single-cell chromosome copy number profiles inferred from scRNA-seq data of K1088N primary PTEC organoid cells after *VHL*-ASK with or without CA9+ flow sorting. Each row represents an individual cell; columns correspond to chromosomes, scaled by the number of expressed genes. Copy number status for the target chromosome is shown in blue (loss), red (gain) or white (neutral). The total number of cells analyzed (n), and the percentage of cells with the indicated aneuploidy status are shown. (H) Scatter plots showing the CA9 and DAPI staining intensities of K1088N PTECs after the indicated treatment. The gating strategy to flow sort live CA9-positive cells (DAPI– and CA9+) is shown. (I) Violin plots showing the signature scores of a curated HIF pathway activation gene set (see Methods) in K1088N PTECs after the indicated treatment. The dots represent individual cells. The dashed line represents the calculated threshold (see Methods) for defining cells with a high (HIF-High) or a low (HIF-Low) HIF pathway signature score, used for assigning *VHL* proficient (*VHL*+) or *VHL* loss (*<u>VHL-</u>*) status.

**Figure S11.**
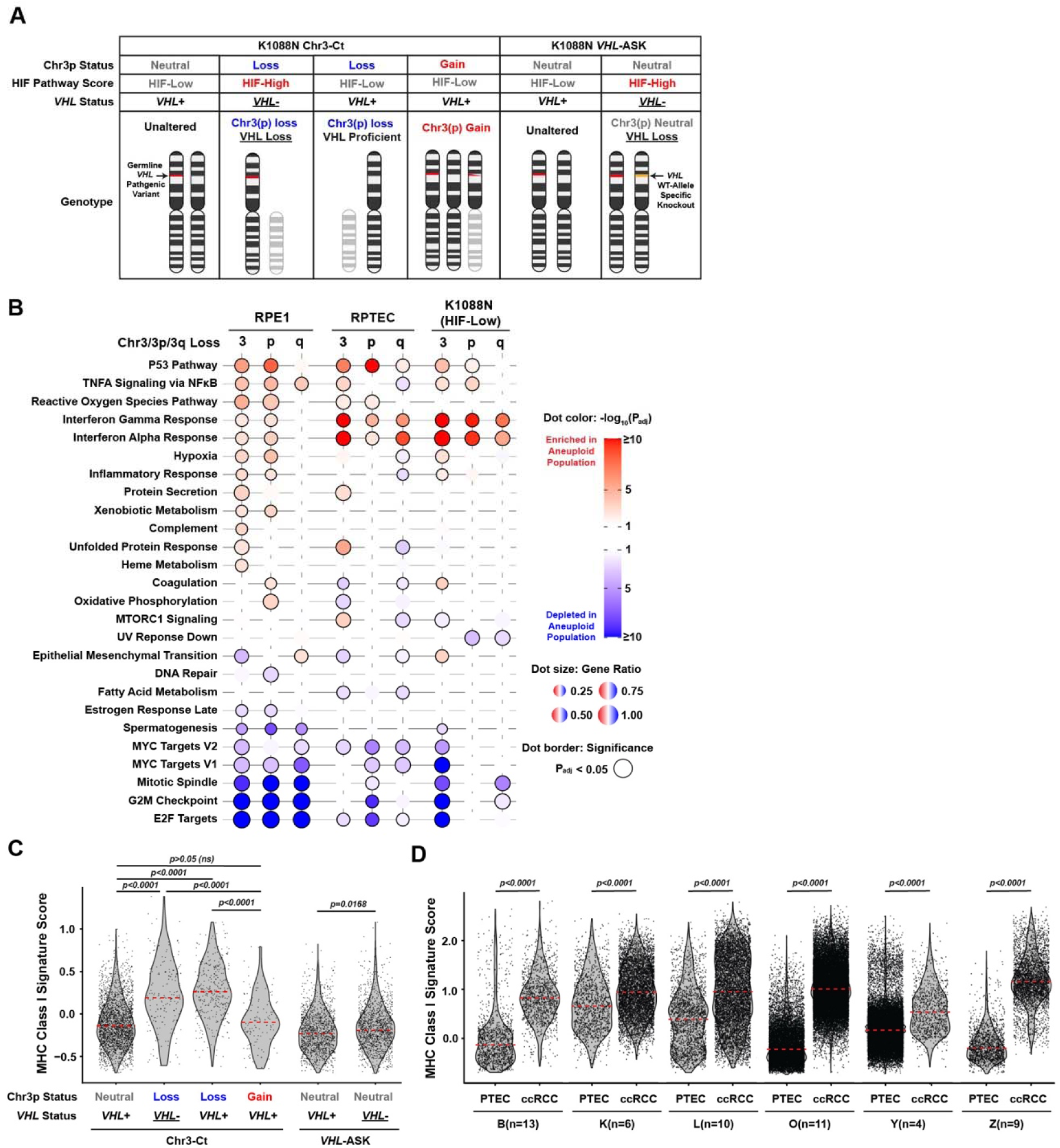
Comparison of transcriptomic alterations associated with Chr3(p) loss in different cell backgrounds, related to Figure 6. (A) Schematics showing the classification of the engineered K1088N cell populations of interest with scRNA-seq data. The classification first considers the treatment (Chr3-Ct or *VHL*-ASK), and subsequently the inferred Chr3(p) status (Neutral, Gain or Loss, Figure 6A) and inferred *VHL* status from the HIF pathway score (Figure S10I). The genotypes of the classified groups are shown with the corresponding Chr3(p) copy number and *VHL* status. (B) Dot plot showing significantly enriched or depleted hallmark gene sets by GSEA in RPE1, RPTEC, and K1088N cells with indicated whole-Chr3 or Chr3-arm losses, compared to the corresponding copy-neutral population. The analyses exclude genes located within the corresponding aneuploid region to highlight secondary genome-wide effects. The columns represent the indicated whole-chr or chr-arm loss. The color gradient represents –log_10_(P_adj_) values with red and blue corresponding to positive and negative normalized enrichment scores, respectively. Dots with a solid border highlight significant changes (P_adj_ <0.05). (C and D) Violin plots showing the signature scores for MHC Class I genes in the engineered K1088N cells with the indicated genotypes (C) and the paired ccRCC and PTEC cells from patient tumor and normal-adjacent samples, respectively, in the published ccRCC tumor scRNA-seq cohorts (B^52^, K^53^, L^54^, O^55^, Y^56^, and Z^57^) (D). Each dot represents an individual cell, and the red dashed line represents the median score (Kruskal-Wallis test followed by Dunn’s post hoc multiple comparison test with adjusted p values shown).

## SUPPLEMENTAL INFORMATION

**Table S1 CRISPR-Taiji gRNA information, related to** Figure 1

**Video S1 Split chromosome arms induced by CRISPRt, related to Figure 3** Representative video clip of time-lapse live-cell imaging showing the induced chromosome mis-segregation in an RPE1 cell after CRISPRt with Chr3-sg2 gRNA. One mis-segregated chromosome presents the phenotype of split chromosome arms into the two daughter cells with a putative centromere bridge, and the other mis-segregated chromosome simply forms a micronucleus in one daughter cell. The left view shows the merged ‘Trans’ channel (cell) in grey and ‘mTurquoise-H2B’ channel (nucleus) in blue. The right view shows only the nucleus channel in grey. Scale Bars, 30 μm.

## EXPERIMENTAL MODEL AND STUDY PARTICIPANT DETAILS

### Human participants

Two male von Hippel-Lindau (VHL) disease patients, designated K985 and K1088, with VHL germline variant c.407T>C (p.F136S) and c.256 C>T (p.P86S), respectively, were recruited to the TRACERx Renal study with patients’ consent at the Barts Health NHS Trust, London, UK. The TRACERx Renal study is an ethically approved prospective cohort study (NCT03226886, National Health Service Research Ethics Committee approval 11/LO/1996) sponsored by the Royal Marsden NHS Foundation Trust and coordinated by its Renal Unit. Peripheral blood mononuclear cells (PBMCs) and renal surgery specimens, including normal kidney specimens, were collected from the recruited patients. Specimens were handled and stored according to standard protocols for downstream workflows.

### Cell lines

The induced pluripotent stem cell (iPSC) line iFCI013 (also known as CRICKi010-A) was reprogrammed from the PBMCs of patient K985 at the Francis Crick Institute. The reprogramming steps and quality validation of iFCI013, including karyotyping and pluripotency marker expression, have been published^70^. iFCI013 cells were maintained in mTeSR Plus medium (STEMCELL Technologies) on cultureware coated with Cultrex ReadyBME (R&D Systems), at 37°C in a humidified incubator with 5% CO2.

The primary renal proximal tubule epithelial cells (PTECs) K1088N were derived from macroscopically normal kidney tissue obtained from patient K1088. The kidney tissue was processed into small fragments and digested with 2 mg/mL collagenase P (Roche, 249002001) at 37°C for 1 h on a shaking platform to release single cells. The digested suspension was filtered through a 40 µm cell strainer to remove undigested tissue fragments, followed by centrifugation to pellet the cells. The cell pellet was resuspended in cold growth factor-reduced Matrigel (Corning, 356231) and plated on pre-warmed cultureware as a 3D organoid culture. Organoids were maintained in kidney tubuloid medium (Advanced DMEM/F12 supplemented with 1x GlutaMAX, 1x B27 supplement, 10 mM HEPES, 1 mM N-acetyl-L-cysteine, 5 µM A83-01, 50 ng/mL EGF, 100 ng/mL FGF-10, 0.1 mg/mL primocin, and 10% R-Spondin 1 conditioned medium) at 37°C in a humidified incubator with 5% CO2^79^.

The RPE1 (*hTERT*-immortalized retinal pigment epithelial cells) cell line, obtained from ATCC, was maintained in DMEM (Sigma, D6429) supplemented with 10% fetal bovine serum (FBS) and 100 U/mL Penicillin-Streptomycin, at 37°C in a humidified incubator with 5% CO2. The RPE1 cell line expressing mTurquoise-H2B for live-cell imaging was generated and characterized in the previous publication^71^.

The RPTEC (*hTERT*-immortalized renal proximal tubule epithelial cells) cell line was obtained from ATCC. The RPTEC-shP53 cell line (expressing a short hairpin RNA against TP53), was generated from the parental RPTEC cell line by retroviral transduction of pSRZ.p53shRNA retroviral construct (a gift from Dr. Jerry Shay’s lab, UTSW) expressing a short hairpin RNA against TP53. The RPTEC and RPTEC-shP53 cells were maintained in renal epithelial cell growth basal medium (Lonza) supplemented with 0.5% tetracycline-free FBS, 100 U/mL Penicillin-Streptomycin, 10 ng/mL hEGF, 1x insulin-transferrin-selenium solution, 1 μg/mL hydrocortisone, 10 μM epinephrine, 50 ng/mL triiodo-l-thyronine, 30[μg/mL gentamicin and 15 ng/mL amphotericin B at 37°C in a humidified incubator with 5% CO2.

## METHOD DETAILS

### dCas9/HiFi-Cas9-gRNA RNP Electroporation

The single-guide RNAs (sgRNAs) used for CRISPRt screening were purchased from Genscript Biotech. The CRISPR RNAs (crRNAs, XT version) and trans-activating crRNA (tracrRNA)-Atto550 conjugate used for immunofluorescence imaging were purchased from IDT. All synthetic RNAs are synthesized with chemical modifications at the first three 5’ and 3’ terminal RNA residues, which protect them from nucleases. All synthetic RNAs were resuspended in low-EDTA TE buffer to a stock concentration of 100 µM. High-fidelity Cas9 (Alt-R S.p. HiFi-Cas9 V3, 10 µg/µL) and catalytically dead Cas9 (Alt-R S.p. dCas9 V3, 10 µg/µL) were purchased from IDT.

For preparing CRISPR ribonucleoprotein (RNP) with sgRNAs for a 20 µL (or 100 µL) electroporation reaction, mix 1 µL (or 3 µL) of 100 µM sgRNA with 0.5 µL (or 1.5 µL) of dCas9 or HiFi-Cas9 in a PCR tube. For preparing RNP with crRNAs for a 20 µL electroporation reaction, mix 0.75 µL of 100 µM crRNA with 0.75 µL of 100 µM tracrRNA-Atto550 in a PCR tube; then anneal the crRNA and tracrRNA by incubating at 95°C for 5 min, followed by a gradual cool down to 25°C with a ramp rate of 0.2°C/s in a thermocycler; then mix the crRNA/tracrRNA complex with 0.5 µL dCas9. Incubate the mixed dCas9/HiFi-Cas9 and gRNA at room temperature for 30 min (in the dark if using crRNA-tracrRNA-Atto550).

The electroporation was performed with the Lonza P3 Primary Cell 4D-Nucleofector X kit S (20 µL reaction) or L (100 µL reaction), with Lonza 4D-Nucleofector X Unit according to the manufacturer’s instructions. Briefly, single-cell suspension was prepared using Accutase treatment for iFCI013 cells, TrypLE treatment for RPE1 cells, and Dispase followed by Accutase treatment for K1088N organoids. For each 20 µL (or 100 µL) electroporation reaction, 200k (or 1000k) cells were transferred to a sterile tube and centrifuged. All the supernatant was carefully removed to avoid negative impacts on electroporation efficiency. The cell pellet was then resuspended in 20 µL (or 100 µL) of complete P3 electroporation buffer, prepared by mixing 16.4 µL (or 82 µL) of P3 Solution with 3.6 µL (or 18 µL) of Supplement 1. The cell suspension was gently mixed with the corresponding RNP complex by pipetting without introducing bubbles and then transferred to the electroporation well or cuvette. Electroporation pulses were immediately applied using the Lonza 4D-Nucleofector X Unit with pulse codes CA-137 (iFCI013 cells), EA-104 (RPE1 cells), or CM-138 (K1088N and RPTEC cells).

After electroporation, 135 µL (or 675 µL) of the corresponding culture medium described below was added to the electroporation well or cuvette with gentle mixing, and the cells were incubated for 5 min. If using sgRNA, the cell mixture was directly seeded into suitable cultureware after gentle mixing. If using rRNA/tracrRNA-Atto550, cells were centrifuged to remove all the supernatant containing the excess fluorescent RNP complex, before resuspending and seeding into the appropriate cultureware. Post-electroporation, iFCI013 cells were cultured on Cultrex ReadyBME-coated cultureware or glass coverslip with mTeSR Plus medium supplemented with CloneR2 (STEMCELL Technologies) to support single-cell survival; K1088N cells were cultured on Matrigel-coated cultureware or glass coverslip with the kidney tubuloid medium supplemented with 1% Matrigel as Matrigel-sandwiched 2D culture (reduced environmental hypoxia compared to 3D culture); RPE1 and RPTEC cells were cultured at the corresponding standard conditions.

### dCas9-gRNA RNP Lipofectamine Transfection

The lipofectamine CRISPRMAX transfection reagent (Invitrogen) was used to reverse-transfect the dCas9-sgRNA RNP complex into RPTEC-shp53 cells. Briefly, mix 1 µL of 100 µM sgRNA, 1.2 µL of dCas9, and 10 µL of Cas9 Plus reagent in 250 µL Opti-MEM in one tube, and mix 15 µL of CRISPRMAX reagent in 250 µL Opti-MEM in another tube. Mix the contents of the two tubes and incubate for 10 min at room temperature. Mix the lipofectamine-RNP mixture with 1.2 million RPTEC-shp53 cells resuspended in 2.5 mL medium before seeding into a 6 cm dish. Replace the medium after 24 h.

### CRISPR-Taiji gRNA Screening and Validation

For each chromosome, the designed gRNAs with more canonical binding sites within the corresponding CENPA-bound region were prioritized for screening in batches until functional gRNAs were identified or all designed gRNAs were tested. The screened gRNAs were assigned a CRISPRt ID, for example, ‘Chr1-sg1’ representing Chr1-specific sgRNA1.

For gRNA screening and validation, the dCas9-sgRNA RNP complex was introduced into iFCI013, RPE1, or K1088N cells using the 20 µL electroporation reaction system with the 16-well electroporation strips as described above. For each batch, the 3 non-targeting (NT) sgRNAs were included as the negative control. For each 16-well electroporation strip, a maximum of 15 candidate sgRNAs were tested, and Chr3-sg2 was included as the positive control to check electroporation efficiency. Following electroporation, for each gRNA, 65k cells were seeded into a 24-well plate with 13mm coverslips, while 135k cells were seeded into a 12-well plate with the corresponding medium described above. Cells were cultured for 48 h before characterization.

For micronucleated cell quantification, coverslips were fixed in 4% paraformaldehyde (PFA). The fixed cells were then permeabilized and stained with PBS containing 0.2% Triton X-100 and 1 µg/mL DAPI for 5 min. DAPI images were captured using the Echo Revolve epifluorescence microscope with a 20x objective. The number of nuclei per image was quantified using CellProfiler software, and the number of micronucleated cells was manually counted in a blinded manner to ensure unbiased assessment.

The screening results are summarized in Table S1. Functional gRNAs that have been extensively validated across multiple cell models are highlighted in red. Additional functional gRNAs, highlighted in orange, demonstrate good micronucleus induction efficiency in the initial screen but have not yet validated across models. While functional gRNAs highlighted in red consistently induce micronucleus formation across the models we have tested, some degrees of model-specific variations in gRNA efficiency were observed. Therefore, we recommend testing multiple functional gRNAs (both red and orange, if available) for the chromosome of interest to identify the most efficient gRNA when establishing CRISPRt in a new cell line. In addition, RNP delivery is the preferred method for efficient and reproducible engineering using CRISPRt.

### Fluorescence *in situ* Hybridization (FISH)

For validating chromosome identity within the micronucleus, single-cell suspensions were generated using TrypLE treatment (important to make sure that the Cultrex/Matrigel coating is fully digested). The cell pellet was gently dissociated by tapping the tube, then fixed by slowly adding ice-cold Carnoy’s fixative (methanol and acetic acid in a 3:1 volume-to-volume ratio) dropwise while gently vortexing. Cells were incubated at room temperature for 20 min. After centrifugation to remove the fixative, the cells were subjected to a second round of fixation with the same procedure. The fixed cell pellet was resuspended in 50 µL of ice-cold Carnoy’s fixative. Cells were subsequently dropped onto slides and air-dried for further processing.

For metaphase spread and chromosome painting, RPTEC-shP53 cells were treated with 120[ng/mL KaryoMAX colcemid for 6 h before collection. Using the RPTEC cells with *TP53* knockdown facilitates the survival and cell cycle progression of aneuploid cells for metaphase spread analysis. Cell pellets were gently resuspended in 5 mL of 75 mM KCl solution dropwise while gently vortexing. Cells were incubated for 6 min in a 37 °C water bath and then fixed by slowly adding ice-cold Carnoy’s fixative, followed by centrifugation and resuspension in Carnoy’s fixative. Cells were subsequently dropped onto slides to form metaphase spreads and air-dried for further processing.

For staining, the slides were then rehydrated in 2x Saline-Sodium Citrate (SSC) buffer for 2 min, followed by sequential dehydration in an ethanol series (70%, 85%, and 100%) for 2 min each. Next, the cell spots were hybridized with diluted centromere probes (OGT) or telomere probes (OGT), corresponding to the target chromosomes (Figure S1B), or with Chr3 chromosome painting probes (MetaSystems) for RPTEC-shP53 metaphase spread staining. Briefly, the probes were applied to the cell spot on the slide and covered with a 13mm coverslip, which was then sealed with rubber solution. Slides were incubated at 75°C for 2 min for denaturation, followed by overnight hybridization in a humidified dark chamber at 37°C. The following day, coverslips were removed, and the slides were immersed without agitation in 0.4x SSC buffer at 72°C for 2 min, and then in 2x SSC buffer with 0.05% Tween-20 at room temperature for 30 s. The slides were counterstained with DAPI, air-dried, and mounted in antifade mounting solution.

Micronucleus FISH images were captured using the Echo Revolve epifluorescence microscope with a 20x objective. Micronuclei with or without the FISH signals of the corresponding target chromosome were manually quantified. Metaphase chromosome painting images were acquired on the Metafer Scanning and Imaging Platform microscope (MetaSystems) with a 63x objective. Metaphase images were analyzed using the Isis Fluorescence Imaging Platform (MetaSystems, v.5.8.15).

### Immunofluorescence and quantification

The dCas9-crRNA-tracrRNA-Atto550 RNP complex was introduced into iFCI013 cells by electroporation as described above. At indicated times post-electroporation, iFCI013 cells seeded on coverslips were fixed using ice-cold methanol for 30 min at –20 °C. After fixation, the cells were rehydrated in PBS for 5 min at room temperature and washed twice with PBS to remove residual methanol. Blocking was performed with 5% milk-PBS for 1 h at room temperature, followed by two PBS washes to remove any remaining blocking solution. Primary antibodies were diluted in 2% BSA-PBS, and the cells were incubated with the primary antibody solution overnight at 4 °C. The primary antibodies used are Anti-Centromere Protein (CREST, 1:300, Antibodies Inc, 15-234), anti-HEC1 (1:300, Invitrogen, MA1-23308), and anti-α-Tubulin (1:300, Invitrogen, A11126). After primary antibody incubation, cells were washed three times in PBS on a shaking platform for 5 min each. Secondary antibody incubation was performed in 2% BSA-PBS for 1 h at room temperature. The secondary antibodies used are Anti-Mouse-IgG-Alexa-488 (1:500, Invitrogen, A332766TR) and Anti-Human-IgG-Alexa-647 (1:500, Invitrogen, A-21445). Following secondary antibody incubation, the cells were washed three times in PBS for 5 min each. Coverslips were then mounted with antifade mounting medium containing DAPI. Z-stack fluorescence images were captured using a Zeiss Invert880 confocal system with Zeiss ZEN software with a 63x oil immersion lens.

Quantification of dCas9 (Atto550-gRNA) recruitment and signal intensity at 8 h post-electroporation was performed using CellProfiler software. First, nuclei were segmented based on the DAPI signal to generate a nucleus mask. CREST foci were identified within the nucleus mask and expanded by 3 pixels to create the CREST foci mask (as CREST and Atto550-gRNA foci may not always co-localize perfectly). Then, Atto550-gRNA foci were then detected within the CREST foci mask and assigned to their corresponding nuclei, enabling per-nucleus quantification of foci number. For intensity analysis, the identified Atto550-gRNA foci were further expanded by 3 pixels to create the Atto550-gRNA foci mask, and integrated signal intensities from both the Atto550-gRNA and CREST channels were measured within this mask. To account for imaging variability, the integrated Atto550-gRNA foci intensity was individually normalized to the corresponding CREST foci intensity.

Quantification of the kinetochore rupture (CREST staining) at the target centromere (Atto550-gRNA foci) at 24 h post-electroporation across cell cycle stages was performed manually. Briefly, the cells were first classified into cell cycle stages based on their nuclear morphology. Then, the target chromosomes with Atto550-gRNA foci were classified into normal and abnormal locations according to the cell cycle stages. Finally, the classified chromosomes with or without CREST foci were quantified.

### ATAC-seq library preparation and sequencing

The dCas9-sgRNA RNP complex was introduced into iFCI013 cells by electroporation as described above, and experiments were performed with two independent biological replicates. Single cells were harvested using TrypLE treatment 8 h post-electroporation, allowing sufficient time for dCas9 binding with minimal mitotic progression. ATAC-seq library preparation followed the published ATAC-seq protocol^80^, with all reagents and buffer recipes following the protocol unless otherwise specified. Briefly, 50k cells were used as input for each sample. Nuclei were prepared by incubating cells in ATAC-seq Lysis Buffer for exactly 3 min on ice. The nuclei were then incubated with the Transposition mix at 37 °C for 30 min in a thermomixer shaking at 1,000 rpm, generating transposed fragments. These fragments were purified and eluted in nuclease-free water using the DNA Clean & Concentrator-5 kit (Zymo). The purified fragments were then barcoded and amplified by PCR, and the exact number of amplification cycles for each sample was determined using the NEBNext Library Quant Kit for Illumina, according to the manufacturer’s protocol. The quality and fragment size of the final libraries were assessed using the TapeStation 4200 (Agilent Technologies). Libraries were sequenced to 25 million PE-100bp target read pairs per sample using the Illumina NovaSeq 6000 at the Genomics Science Technology Platform at the Francis Crick Institute.

### Time-lapse Live-cell imaging

The dCas9-sgRNA RNP complex was introduced into RPE1 cells expressing mTurquoise-H2B by electroporation as described above. After electroporation, 30k cells per well were seeded into an 8-well chambered polymer coverslip (Ibidi, 80806) and allowed to settle for 16 h. Time-lapse live-cell imaging was then performed using a Nikon Eclipse Ti inverted microscope, equipped with a custom humidified enclosure (Okolabs) that maintains 37°C with 5% CO2. The Nikon Perfect Focus System (PFS) was used for auto-focus during image acquisition. Phase-contrast and fluorescent images were captured every 3 min for 48 h using ImageJ-mManager software with a 40x objective. Laser intensity and exposure time were optimized to prevent phototoxicity or photobleaching in the cells. Image processing was performed using FIJI software, and chromosome mis-segregation events were analyzed and quantified manually. The mitotic event was first classified by the presence of micronucleus formation, and then mitotic events with micronucleus formation were further classified by the presence of chromosome arm split (Video S1). The mitotic duration was quantified as the time interval between the start of chromosome compaction in prophase and the end of chromosome decompaction in telophase. The statistical significance was calculated with a two-tailed Mann-Whitney test.

### *VHL* Allele-specific Knockout

The gRNA spacer VHL-sg (5’-CATACCGAGTTGAAGCTGCC –3’) was designed to specifically target the wild-type VHL allele while sparing the germline variant allele of patient K1088. This selective targeting was achieved by designing VHL-sg adjacent to the VHL germline variant of patient K1088; therefore, the SpCas9 protospacer-adjacent motif (PAM) site (5’-NGG-3’), essential for Cas9 binding, is altered from 5’-GGG-3’ to 5’-GGA-3’ by the germline variant, which prevents Cas9 from targeting the germline variant allele. The high-fidelity Cas9 (HiFi-Cas9) was used to further ensure the allele specificity. The HiFi-Cas9-sgRNA RNP complex was introduced into K1088N cells via electroporation as described. To confirm the successful introduction of frameshift mutations, Sanger sequencing was performed 11 days post-electroporation. The primer sequences used for PCR amplification of VHL exon 1 are VHLe1-FW (5’-TGGTCTGGATCGCGGAG –3’) and VHLe1-RV (5’-CTTCAGACCGTGCTATCGTC –3’).

### Flow Cytometry and Cell Sorting

The dCas9-sgRNA RNP complex was introduced into K1088N cells by electroporation as described above. Cells were initially cultured for 7 days to confluence before splitting at a 1:6 ratio, and the cells were further cultured for 4 days before harvesting for flow sorting. Single-cell suspension was generated with Accutase treatment, and the cell pellet was washed with PBS once. Cells were stained with 1:50 diluted anti-CA9-PE antibody (Miltenyi, 130-123-299) in flow sorting buffer (1% bovine serum albumin (BSA) and 1μM EDTA in PBS) for 30 min at 4 °C in the dark. The cell pellet was washed with flow sorting buffer twice before being resuspended in 250 μl flow sorting buffer for every 1 million cells supplemented with 1 μg/mL DAPI. Cell sorting was performed on a BD FACSAria Fusion cell sorter with BD FACSDiva software. CA9 and DAPI gate thresholds were determined with the non-targeting dCas9-NT-sg1 (CTRL) sample. Singlet cells that are CA9 positive and DAPI negative for HiFi-Cas9-VHL-sg (*VHL*-ASK) or dCas9-Chr3-sg2 (Chr3-Ct) samples were sorted with ‘Yield’ precision mode scRNA-seq characterization.

### scRNA-seq Library Preparation and Sequencing

The peak of chromosome mis-segregation occurs between 24-48 h post-electroporation. Early time points allow better capture of aneuploid cells, particularly for gRNA with low mis-segregation induction efficiency, as aneuploidy in non-transformed cells typically confers a fitness disadvantage relative to parental cells. Conversely, later time points are more suited for detecting secondary genome-wide effects of aneuploidy, due to the temporal delay from transcriptomic alterations to proteomic changes that mediate downstream phenotypes. IFCI013 cells were harvested as single-cell suspension at an earlier time point of 72 h following CRISPRt to demonstrate chromosome-specific aneuploidy generation. RPE1 and RPTEC cells were harvested as single-cell suspension at a later time point of 120 h (5 days) following CRISPRt to allow additional 72-96 h for the establishment of secondary genome-wide effects (72 h is mostly sufficient to have the secondary genome-wide effect in genetic perturbation experiments, which is also validated later by our transcriptomic analyses of populations with Chr21 gain or loss. K1088N cells were harvested as single-cell suspension at later time points of 7 or 11 days with or without CA9 flow-sorting enrichment following CRISPRt to focus on the secondary genome-wide effect (these time points reflect the durations required to reach confluence following seeding or passaging).

For single-cell fixation, up to 1 million singlet cells from each sample were fixed using the Parse Bioscience Evercode Fixation v3 kit, then cryopreserved as aliquots at –80 °C following the manufacturer’s protocol. After fixation, single-cell barcoded cDNA libraries were prepared using the Parse Bioscience Evercode WT (or WT Mini) v3 kit, targeting 1,650 cells per sample according to the manufacturer’s instructions. The quality and fragment size of the final library (consisting of two sub-libraries) were assessed using the TapeStation 4200 (Agilent Technologies). The library was sequenced to an average depth of 100k PE-100bp read pairs per cell using the Illumina NovaSeq 6000 at the Genomics Science Technology Platform at the Francis Crick Institute.

## QUANTIFICATION AND STATISTICAL ANALYSIS

### Statistics and reproducibility

The number of biological replicates and the minimum number of quantified events are indicated in the corresponding figure legends. Error bars represent the standard deviation (SD). Statistical tests are detailed in the respective figure legends unless they are part of the default analysis package settings. For multiple hypothesis testing, adjusted p-values were calculated using the Benjamini– Hochberg procedure.

### CRISPR-Taiji gRNA Design

The design and annotation steps of CRISPR-Taiji (CRISPRt) guide RNA (gRNA) spacer sequences are outlined in Figure S1A with gRNA spacer details summarized in Table S1.

The complete “Telomere-to-Telomere” (T2T) reconstruction of a human genome with Y (T2T-CHM13v2.0) and centromere/satellite repeat annotations (Cen/Sat v2.0) from the T2T Consortium, was downloaded from <u>github.com/marbl/CHM13</u>^16–18^. For this study, Cen/Sat v2.0 annotation was used to select the active α-satellite higher-order-repeat region (active αSat HOR) and the entire (peri)centromere regions (pan-centromere). Additionally, CENPA Cut&Run characterization data for the T2T-CHM13v2.0 assembly, generated by the T2T Consortium, was used to select the CENPA-bound regions^17,18^.

The crisprDesign R package was used to identify all Streptococcus pyogenes Cas9 (SpCas9) gRNA spacers within the CENPA-bound sequences for each chromosome^72^. All gRNA spacers were subsequently filtered based on the following criteria: GC-content between 35% and 80%, and a minimum of 20 canonical binding sites within the CENPA-bound region of at least one human chromosome. The GC-content range was selected based on previously published findings^81^. The CENPA-bound chromosome specificity was calculated as the number of binding sites in the CENPA-bound region of a specific chromosome divided by the total number of binding sites in the CENPA-bound regions across all chromosomes. A specificity threshold of ≥0.9 was applied, resulting in 1,830 chromosome-specific gRNAs across 24 human chromosomes. In addition, 15 bi-chromosome-specific gRNAs with binding sites on both Chr13 and Chr21 (i.e. not satisfying ≥0.9 CENPA-bound chromosome specificity) were included for efficiency improvement. Finally, 3 non-targeting gRNAs were included as negative controls.

Further analyses were conducted to calculate the chromosome specificity of the CRISPRt gRNA set across the active α-sat HOR region and the pan-centromere region (similar to CENPA-bound chromosome specificity but across different centromeric regions). gRNAs with active αSat HOR or pan-centromere chromosome specificity <0.9 were flagged in red but not excluded (Table S1). Additionally, the centromere specificity of the gRNAs was calculated as the total number of binding sites across the pan-centromere region divided by the total number of binding sites across the entire genome (T2T-CHM13v2.0). All selected CRISPRt gRNAs demonstrated a centromere specificity score >0.98, indicating limited on-target binding sites of CRISPRt gRNAs outside the centromeric region.

### ATAC-seq analysis

The ATAC-seq library was first processed using the nf-core/atacseq pipeline (https://nf-co.re/atacseq/2.1.2) for quality checks^73^. To assess the centromeric ATAC-seq signals, we adapted the previously published method for CENPA CUT&RUN data analysis with minor modifications^17^. Briefly, ATAC-seq raw reads were processed with TrimGalore (v0.6.1) to remove the adapter sequence. Trimmed reads were aligned to the T2T-CHM13v2.0 reference or a customized reference consisting of the CENPA-bound sequences used for gRNA design (T2T-CENPA) using BWA (v0.7.17) with the parameters: bwa mem –k 50 –c 1000000 [reference] [R1] [R2]^17,74^. The resulting sam files were filtered using SAMtools (v1.13) with the command: samtools view –b –F 2308 [sam], which randomly assigns reads with multi-mapping sites to a single mapping location to prevent mapping biases in highly repetitive regions. PCR duplicates were marked and removed using Picard (v2.25.1). For visualization, the processed BAM files were converted to bigwig files using deepTools (v3.5.1) and normalized by the number of mapped reads.

To quantify chromosome-specific changes in ATAC-seq signals, we calculate the percentage of reads mapped to the entire chromosome (with T2T-CHM13v2.0 reference) or the CENPA-bound region (with T2T-CENPA reference) for individual chromosomes. Average log2 fold changes (log2FC) were then calculated by comparing to NT-sg1. Due to the repetitive nature of centromeric sequences, it remains challenging to accurately map centromeric reads and the mapped reads to the T2T-CENPA reference may derive from broader αSat regions. In addition, we assumed that CENPA-bound regions in iFCI013 cells are identical to the published T2T reference, which may not be completely accurate. Therefore, we noted the baseline distribution of the reads mapped to the T2T-CENPA reference may not reflect the ground truth, however, the relative accessibility changes compared to NT-sg1 remain reliable.

### Parse scRNA-seq Analysis

For Parse scRNA-seq, raw FASTQ reads were processed using the proprietary Parse Bioscience pipeline (v1.2.1) with default parameters to generate the filtered raw cell-gene count matrix. The filtered matrix was then used to generate the Seurat objects, excluding features in fewer than 3 cells and cells with fewer than 200 features using the Seurat package (v5.0.0)^75^. For the filtering steps, cells with more than 20% mitochondrial reads were excluded as dead cells, and cells with feature counts more than the mean plus 2x standard deviation were excluded as potential doublets. In addition, “MALAT1” and mitochondrial genes were removed for the downstream analyses. Normalization and SCTransform functions were applied, followed by UMAP dimensionality reduction and clustering using the Seurat package^75^. Single-cell copy number inference from scRNA-seq data was performed using the InferCNV package (v1.21.0)^76^. Briefly, residual gene expression was calculated and denoised using the NT-sg1 (CTRL) sample as the unaltered ‘copy-neutral’ reference. Copy number heatmaps were then generated by plotting the residual gene expression across each chromosome (scaled by the number of genes included in the InferCNV calculation). To assign cell populations with different aneuploidy statuses, average residual gene expressions across chromosome arms were calculated. Thresholds for defining copy number loss or gain were set at the average residual gene expressions, where 1% of NT-sg1 cells were classified as loss or gain, respectively. Cells between these two thresholds were classified as copy number neutral. The assigned aneuploidy population indeed reflect expression changes across the whole chromosome or the entire chromosome arm until the centromere location.

For allele-specific expression analysis in RPE1 and RPTEC cells, BAM files generated by the Parse Bioscience pipeline were demultiplexed into individual BAM files based on the CRISPRt treatment (sample barcode) and aneuploidy status (cell barcode). For each CRISPRt treatment, variants were called on the corresponding target chromosome(s) using bcftools (v1.9) in multiallelic mode, based on reads from the unaltered (copy number neutral) population. SNPs with a quality score >30 and read depth >20 were retained. Allele-specific expression was quantified using GATK ASEReadCounter (v4.4.0.0) for both unaltered and whole-chr loss populations. SNPs with total read depth >20 in both populations were considered high-confident. The average alternate allele fraction of these high-confident SNPs was calculated and visualized for each target chromosome.

For transcriptomic analyses of each assigned aneuploid population in RPE1 and RPTEC cells, differentially expressed genes (DEG) for the classified aneuploid population compared to the corresponding unaltered ‘copy-neutral’ population from the same CRISPRt treatment were identified using the FindMarkers function of the Seurat package^75^. We noted that minor aneuploid populations with concurrent loss of one arm and gain of the other were also generated by CRISPRt, which were excluded from the current analysis of chr-arm aneuploidies. To highlight the secondary genome-wide effect, genes on the corresponding affected aneuploidy region were filtered out from the DEG list for subsequent analyses unless specified. Gene set enrichment analysis (GSEA) was then performed with the filtered DEG output for Human MSigDB hallmark gene sets and a curated hypoxia-inducible factor (HIF) pathway activation gene set (Lombardi HIF)^37^ using the fgsea package (v1.24.0)^77,78,82^. Gene sets with an adjusted p-value < 0.01 in at least one aneuploid group were selected for visualization.

For gene set alteration co-occurrence analysis, the number of times when a pair of hallmark gene sets were both significantly altered in the same whole-chr aneuploid population was quantified as the total co-occurrence count. The directionality bias was calculated as the difference between times when both are significantly enriched or depleted (same direction) and times when one is significantly enriched and one is significantly depleted (opposite directions). The co-occurrence directionality score was then calculated as the directionality bias divided by the total co-occurrences count, yielding a score that ranges from –1 (always opposite directions) to 1 (always same direction), with 0 indicating no directional consistency. Statistical significance of the observed directionality was assessed using a two-sided binomial test, testing against a null hypothesis of no directional bias (i.e., expected score of 0). Multiple testing correction was performed using the Benjamini–Hochberg procedure. For quantifying the overlaps of the leading-edge gene subsets between a pair of hallmark gene sets, the Jaccard similarity index was calculated. Statistical significance of the leading-edge gene subset overlaps was assessed using Fisher’s exact test. Multiple testing correction was performed using the Benjamini–Hochberg procedure.

For extracting consistently expressed ribosomal large subunit (RPL) and small subunit (RPS) genes, the percentage of cells with non-zero read count for each ribosomal gene was calculated with scRNA-seq data of RPE1 and RPTEC NT-sg1 (CTRL) treatment. Genes with non-zero read count in more than 50% of cells in both cell lines are defined as consistently expressed RPL and RPS genes.

For transcriptomic analyses in K1088N cells, a *VHL* loss status was also assigned to individual cells based on the Lombardi HIF gene signature score^37^. Briefly, a threshold was set at the score where 1% of CTRL cells were classified as HIF pathway signature high (HIF-High). Cells with scores above this threshold were classified as HIF-High and considered as *VHL* loss (*<u>VHL</u>*<u>-</u>). Cells with scores below this threshold were classified as HIF pathway signature low (HIF-Low) and considered as *VHL* proficient (*VHL*+, with functional *VHL* gene copy). Then, the DEG and GSEA analyses were performed as described above for the indicated cell populations with specific Chr3 aneuploidy status and *VHL* loss status, compared to the unaltered ‘copy-neutral’ population. In addition, the MHC Class I (*B2M*, *HLA-A*, *HLA-B*, *HLA-C*, *HLA-E*, and *HLA-G*) gene signature score was calculated using the Seurat package^75^.

### Published Perturb-seq data analysis

Published normalized pseudo-bulk expression matrices from CRISPRi-perturbation scRNA-seq screens (RPE1 essential-wide and K562 genome-wide)^29^ were downloaded from the Figshare+ website and converted into Seurat objects using the Seurat package^75^. For each perturbed gene, hallmark interferon gamma response signature scores were calculated and plotted against CRISPRi knockdown efficiency, quantified as residual gene expression. Additionally, published CRISPRi screening result (gRNA fold change and significance) for interferon-gamma production in primary CD8⁺ T cells^44^ were retrieved from the supplementary table and used to reproduce volcano plots of perturbation effects.

### Published ccRCC tumor scRNA-seq data analysis

Processed matrix files from six published renal cell carcinoma scRNA-seq cohorts (B^52^, K^53^, L^54^, O^55^, Y^56^, and Z^57^) were downloaded. Seurat objects were generated with only ccRCC tumor and normal adjacent samples included using the Seurat package^75^ and then filtered with the same steps used in Parse scRNA-seq data analysis. Normalization and SCTransform functions were applied, followed by UMAP dimensionality reduction and clustering using the Seurat package^75^.

The cluster of normal PTEC cells was assigned using PTEC marker genes *CUBN* and *PDZK1IP1*. To identify ccRCC cells, we first removed the immune cell clusters using the immune cell marker gene *PTPRC*, and then inferred the chromosome copy numbers of the remaining clusters using the InferCNV package^76^ with the assigned normal PTEC cells as the reference. As Chr3p21 loss and *VHL* loss are the near-universal genetic alterations in ccRCC, we first assigned cells with inferred Chr3p21 loss as ccRCC cells and then confirmed their highly expressed HIF pathway signature gene set. Overall, 1127, 5406, 6606, 27320, 953 and 3305 ccRCC cells from tumors with 1189, 1216, 1989, 12289, 28104 and 1515 PTEC cells from paired normal adjacent tissues were identified in cohort B^52^, K^53^, L^54^, O^55^, Y^56^, and Z^57^, respectively. Then, DEG, GSEA and MHC Class I gene signature score analyses were performed as described above.

## RESOURCE AVAILABILITY

### Lead contact

Requests for further information and resources should be directed to and will be fulfilled by the lead contact, Samra Turajlic.

### Materials availability

All the reagents used in this article are commercially available. The cell models used in the article are available from the corresponding authors upon request, subject to a legal agreement.

### Data and code availability

The scRNA-seq and ATAC-seq data have been deposited in the Gene Expression Omnibus (GEO) with accession GSE278125. The published ccRCC scRNA-seq datasets used in this article are available according to the original publication.

The codes with software versions used for CRISPRt gRNA design, ATAC-seq analyses, immunofluorescence foci quantification, and scRNA-seq analyses are available at https://github.com/FrancisCrickInstitute/CRISPR-Taiji.

Any additional information required to reanalyze the data reported in this paper is available from the lead contact upon request.

## ACKNOWLEDGMENTS

We are grateful to the patients and their families who generously participated in the TRACERx Renal Study. We thank Dr Frank Uhlmann, Dr Greg Findlay, Dr James Turner, Dr Michael Howell, and Dr Maxim Molodtsov from the Francis Crick Institute, London, UK, as well as Dr Francisco Barriga from the Vall d’Hebron Institute of Oncology for the fruitful discussion and valuable feedback to drive the project forward. We thank Dr Siniša Volarević from University of Rijeka, Croatia, for his valuable insights on ribosomopathies. We thank Dr Jerry Shay from UTSW for providing pSRZ.p53shRNA construct. We thank Ms. Laila Parvanta and Ms. Chetna Varsani from Barts Health NHS Trust, London, for their support in recruiting VHL patients and providing surgical specimens. We thank the support from the Science Technology Platforms at the Francis Crick Institute, London, UK, especially the Genomics Science Technology Platform, Flow Cytometry Facility, and Light Microscopy Facility. BioRender icons were used in figure preparation under the Francis Crick Institute’s subscription license. This work was supported by the Francis Crick Institute which receives its core funding from Cancer Research UK, the UK Medical Research Council and the Wellcome Trust (CC2044, S.T.), by Cancer Research UK (C50947/A29911, S.T.), by VHL Alliance (Research Grant 2023, S.T., H.F., D.D., and S.T.C.S.), by US Department of Defense (W81XWH2210764, P.L.), and by Beijing Natural Science Foundation (7242083, L.W.).

## AUTHOR CONTRIBUTIONS

Conceptualization: S.T. and H.F.; Methodology: H.F., L.W., H.J., J.A.D., S.E.M., J.F.X.D., P.L. and S.T.; Investigation: H.F., D.D., R.D., L.W., J.Z., F.B., S.T.C.S., J.W., S.C.J., A.F., M.N., Y.Z., R.W., W.A., O.B., Y.D., Yih. X., W.S., Z.S., H.W. and J.Z.; Resources: S.T.C.S., A.L.C., Z.T., T.B., L.G.D., L.H., E.S.L., S.M.O., S.A. and W.M.D.; Data Curation: H.F., D.D., R.D., L.W., J.Z., B.J.Y.T. and Yim. X.; Formal Analysis: H.F., R.D., B.J.Y.T., A.H. and K.A.L.; Manuscript Preparation: H.F. and S.T. with contributions from all the authors

